# Navigating Human Astrocyte Differentiation: Direct and Rapid one-step Differentiation of Induced Pluripotent Stem Cells to Functional Astrocytes Supporting Neuronal Network development

**DOI:** 10.1101/2024.03.27.586938

**Authors:** Imke M.E. Schuurmans, Annika Mordelt, Katrin Linda, Sofia Puvogel, Denise Duineveld, Marina P. Hommersom, Lisa Rahm, Emma Dyke, Gijs-Jan Scholten, Caroline Knorz, Astrid Oudakker, Hans van Bokhoven, Lot D. de Witte, Clara D.M. van Karnebeek, Alejandro Garanto, Nael Nadif Kasri

**Affiliations:** Radboud university medical center, Amalia Children’s Hospital, Department of Pediatrics, Nijmegen, The Netherlands; Emma Center for Personalized Medicine, Departments of Pediatrics and Human Genetics, Amsterdam University Medical Center, Amsterdam, The Netherlands; Radboud university medical center, Department of Human Genetics, Nijmegen, The Netherlands; Donders Institute for Brain, Cognition and Behaviour, Radboud university medical center, Nijmegen, the Netherlands; VIB Center for Brain & Disease Research, Leuven, Belgium; KU Leuven, Department of Neurosciences, Leuven Brain Institute, Leuven, Belgium; Radboud university medical center, Department of Psychiatry, Nijmegen, The Netherlands; United for Metabolic Diseases, Amsterdam, The Netherlands

## Abstract

Astrocytes play a pivotal role in neuronal network development. Despite the well-known role of astrocytes in the pathophysiology of neurologic disorders, the utilization of human induced pluripotent stem cell (hiPSC)-derived astrocytes in neuronal networks remains limited. Here, we present a streamlined one-step protocol for the differentiation of hiPSCs directly into functional astrocytes without the need for ectopic gene expression or neural progenitor cell generation. We found that culturing hiPSCs directly in commercial astrocyte medium, was sufficient to differentiate hiPSCs into functional astrocytes within five weeks. Validation to varying extents across thirty hiPSC-lines demonstrated consistent astrocyte differentiation with minimal batch-to-batch variability. We confirmed astrocyte identity and functionality of the hiPSC-astrocyte monocultures by immunofluorescence, flowcytometry, RNA sequencing, glutamate uptake assays and calcium signaling recordings. Optimization of the protocol enabled co-culture of hiPSC-astrocytes with Ngn2 hiPSC-derived neurons (iNeurons), promoting neuronal differentiation and synapse formation. Lastly, we used single-cell electrophysiology and multi-electrode arrays to confirm robust neuronal network development in 5-week-old hiPSC-astrocyte and iNeuron co-cultures. This protocol offers a rapid and efficient method to establish all-human astrocyte-neuron co-cultures, facilitating the investigation of cell-type-specific contributions to disease pathogenesis. While validated across multiple hiPSC lines, we actively encourage researchers to test and provide feedback on this protocol to enhance its validation for future iterations.

## Introduction

Glia cells were originally thought to only provide physical and structural support to the neurons, and therefore were referred to as the glue of the brain^1^. Astrocytes are the most abundant glial cell type in the brain. Even though structural support of astrocytes to neurons is still considered essential, over the years, there has been increasing recognition of the functional relevance of the astrocytes in the brain^2^. Nowadays astrocytes are known to support neurons at many different levels: neurogenesis, synaptogenesis including both formation and maturation of the synapses, metabolic support, synaptic plasticity, neurotransmission including uptake of neurotransmitters, as well as support of the blood-brain-barrier and regulation of the blood flow^3–6^. The involvement of astrocytes in the pathogenesis of neurodevelopmental and neurodegenerative disorders is becoming increasingly recognized^7–10^.

Human induced pluripotent stem cells (hiPSCs) have paved the road to study human brain development in a dish^11^, also allowing the use of patient-derived material. One of the frequently used protocols to differentiate hiPSCs into glutamatergic neurons make use of *Ngn2* overexpression, also known as induced neurons (iNeurons)^12–14^. To ensure survival and proper maturation of the iNeurons, co-culturing with functional astrocytes is required, for which often primary embryonic rodent astrocytes are used^12,14^. The use of rodent astrocytes in these co-cultures has significant limitations including genomic and functional differences^15^. Additionally, often wildtype rodent astrocytes are used, which potentially bias the development of diseased neurons. Healthy astrocytes could potentially buffer neuronal effects, especially in the context of metabolic disorders. For these reasons iNeurons should preferentially be co-cultured with hiPSC-derived astrocytes, which will not only circumvent the above-mentioned limitations, but also allows to study the cell-type specific effect of the disease, especially for disorders in which the affected genes are enriched in astrocytes.

Many different hiPSC-astrocyte differentiation protocols have been described in literature^8,16–28^. Generally these protocols can be categorized into two main approaches: differentiation using gene overexpression of transcription factor(s) such as *SOX9*, *NFIA* and *NFIB*^25,26,28^, or differentiation using an intermediate differentiation step towards neural progenitor cells (NPCs)^16,17,27^. Direct differentiation using gene overexpression often results in homogenous cultures of astrocytes^25,26,28^. In line with the large scale of astrocytic functions, increasing evidence shows a wide molecular, morphological, and functional astrocytic diversity *in vivo*^29–31^, which is considered to also be important for *in vitro* neuronal cultures^32,33^. Astrocytes can also be derived from NPCs. This protocol has been shown to result in more heterogeneous astrocyte populations. However, these protocols are often more time-consuming and not always result in pure astrocyte populations^25,26,28^. Overall, functional readouts to validate new protocols to create hiPSC-derived astrocytes are mostly focused on only astrocyte characterization in monoculture, but their ability to support neurons in co-culture is often not or very poorly described.

Given the need for hiPSC-derived astrocytes that can be used for astrocyte disease modelling and support iNeurons when co-cultured together, we aimed to develop an efficient and robust differentiation protocol. We were inspired by the principle “we are what we eat”, previously described by Tcw et al. where they applied primary astrocytic media on NPCs to induce an astrocytic phenotype^17,27^. In line with this principle, we have developed a fast and simple one-step protocol to create functional hiPSC-astrocytes. In this protocol, hiPSCs are directly cultured in commercial astrocyte medium, resulting in functional hiPSC-astrocytes within only five weeks of differentiation, without the use of forced overexpression (∼one month of stable expression of constructs) or an NPC intermediate step (∼one month of NPC differentiation). The astrocyte differentiation trajectory for each hiPSC-line across different batches is highly consistent demonstrating minimal batch-to-batch variability. Identity and functionality of the hiPSC-astrocytes was confirmed using a variety of assays, including RNA sequencing, glutamate uptake assays and calcium signaling recordings. We aimed to develop a robust and reliable protocol to co-culture the hiPSC-astrocytes together with the iNeurons. After extensive testing of various adjusted co-culture protocols, the hiPSC-astrocyte and iNeuron co-cultures exhibited consistent neuronal network development when assessed with multi-electrode arrays (MEAs). Our manuscript offers a thorough, step-by-step description of the optimization process, including negative data.

To further optimize these protocols, we would like to encourage the research community to test these protocols within their own labs using their own hiPSC-lines. We kindly ask to share the data with us and provide (both positive and negative feedback), which will be integrated into the 2.0 version of this manuscript, which aims to validate and improve the protocols described here.

## Methods

### hiPSC cell lines and maintenance

In this study, hiPSC astrocyte differentiation was performed for thirty different hiPSC-lines, which includes commercial control lines as well as patient-derived and isogenic lines (Supplementary Table S1)^34–36^. A subset of these lines has been used for detailed validation of the protocol, which is described in this study. hiPSCs were cultured in Essential 8^TM^ Flex Basal Medium (Gibco; A2858501) supplemented with Primocin (0.1 μg/ml; Invivogen; ANT-PM-2) on Geltrex-coated (Gibco; A1413301) 6-well plates (Corning; 353046) at 37 °C/5% CO_2_. At around 80% confluency the hiPSCs were passaged with ReLeSR (Stemcell Technologies; 100-0483) upon washing in Dulbecco’s phosphate-buffered saline (dPBS; Thermo Fisher; 14190169).

### hiPSC-astrocyte differentiation

Prior to plating of the hiPSCs, 6-well plates were coated with 20 µg/mL Human recombinant laminin-521 (Biolaminin; Biolamina; LN521-05) in 1x dPBS^++^ (Gibco; 14040117) and incubated overnight at 4 °C. To start astrocyte differentiation, hiPSCs were washed with dPBS and subsequently dissociated by incubating with TrypLE (Gibco; 12604021) for 3-5 min at 37 °C/5% CO_2_. Afterwards the hiPSC were collected and washed by spinning down in DMEM/F12 (Gibco; 11320074). Upon removal of the supernatant, the pellet was resuspended in Astrocyte medium (AM; ScienCell®; 1801) supplemented with RevitaCell™ (Gibco; A2644501) and Primocin. Accordingly, 150.000 cells were plated per well on a pre-coated 6-well plate and cultured at 37 °C/5% CO_2_. The next day AM supplemented with Primocin was refreshed to withdraw RevitaCell™ from the medium and to remove dead cells. AM supplemented with Primocin was fully refreshed every other day (e.g., three times a week). At 100% confluency, after around four to five days, the astrocytes were split by dissociation using TrypLE and washed by spinning down in DMEM/F12. The pellet was resuspended in AM supplemented with RevitaCell™ and Primocin and all cells were transferred to a T25 flask (Corning; 430372). The next day AM supplemented with Primocin was fully refreshed to withdraw RevitaCell™ from the medium. When the astrocytes reached again 90-100% confluency the cells were split as described previously, and were completely transferred to a T75 flask (Corning; 430641U) followed by full change of AM supplemented with Primocin the day after. Hereafter astrocytes were split at 90-100% confluency at 1:3 onto <u>new</u> T75 flasks and cultured in AM supplemented with Primocin in the absence of RevitaCell™. After around five weeks of differentiation, medium changes were downscaled to twice a week, as the astrocytes mature and therefore the proliferation rate decreased. Throughout the differentiation process the cells were imaged using the Invitrogen™ EVOS™ XL Core Configured Cell Imager. An additional overview of all materials required for the astrocyte differentiation is provided in Supplementary Table S2.

The protocol described above applies to the majority of the hiPSC-lines used in this study. However, for a subset of hiPSC-lines adjustments to the differentiation protocol had to be made to improve astrocyte differentiation, which are described below. In addition, important notes are described, which have been shown to greatly improve astrocyte differentiation and decrease batch-to-batch variability, and therefore are highly recommended to take into account.

**NOTE A**: To ensure full recovery upon thawing, hiPSCs need to be cultured for at least two weeks before starting astrocyte differentiation.

**NOTE B**: The number of hiPSCs plated on day 0 (days in vitro, DIV 0) may vary from 100.000-300.000 cells per well of a 6-well plate, depending on the proliferation rate of that specific hiPSC-line. For example, if a line is growing very slow e.g., it needs to be split less than once a week (usually every 4-5 days), a total of 300.000 hiPSCs per well would be recommended. In contrast, if a hiPSC line shows a high proliferation rate, e.g., has to be split twice a week, a total of 100.000 hiPSCs per well is recommended. This is to prevent splitting within the first four days of differentiation, which we find to be less optimal for the astrocyte differentiation process.

The number of hiPSCs to plate on DIV 0 should be optimized for each hiPSC-line. In case of a “too low” number of hiPSCs at DIV 0, the astrocytes might stop proliferating due to insufficient cell-cell contact. In contrast, a “too high” number of hiPSCs at DIV 0 might result in a too high astrocyte density at DIV 5, often resulting in increased cell death upon splitting due to insufficient astrocyte differentiation (Supplementary Figure 1A).

**NOTE C**: The standard procedure should be to split with RevitaCell™ during the first ∼three weeks of differentiation and to change medium the day after. After three weeks, the differentiating astrocytes can be split without RevitaCell™. However, if the astrocytes have a very low proliferation rate (e.g., have to be split less than once a week), RevitaCell™ treatment can be prolonged until week five or six, or for the entire differentiation period if needed. If the astrocytes grow very fast (e.g., have to be split twice a week), RevitaCell™ treatment can be stopped by week two.

**NOTE D**: In case the astrocytes have not yet reached 100% confluency, but the cells do not seem to spread out but instead grow on top of each other, the astrocytes can be split 1:1 or 1:2 (Supplementary Figure 1B) in a <u>new</u> flask (Supplementary Figure 1C). This can also be done in case the astrocytes have completely stopped proliferating (Supplementary Figure 1D).

**NOTE E**: Some lines show a very rapid increase in size and drastic decrease in proliferation rate (Supplementary Figure 1D), within the first week of astrocyte differentiation. From the thirty hiPSC-lines, the astrocytes derived from two hiPSC-lines do not seem to proliferate at all upon plating in AM at DIV 0 or quickly thereafter. Therefore, upscaling of the number of astrocytes is limited or even impossible. In addition, due to their relative fast increase in astrocyte volume and decreased proliferation rate, these cells are assumed to be more sensitive to the splitting procedure compared to the other astrocyte cultures, as increased cell death is observed upon splitting of these lines. In order to optimize the astrocyte differentiation for these hiPSC lines, several adjustments have been tested. The following adjustments are currently considered most optimal to improve astrocyte differentiation and survival:

- In order to ensure sufficient cell-cell contact, which promotes astrocyte proliferation, a higher number of hiPSCs can be plated at DIV 0 (300.000 – 500.000 hiPSCs per well).
- The hiPSC cultures of the respective lines also show decreased proliferation (e.g., splitting the hiPSC line once every two weeks). Therefore, hiPSC culturing on human Biolaminin-coated plates instead of Geltrex-coated plates (two weeks prior to the start of astrocyte differentiation), has been shown to enhance the proliferation rate of these hiPSC lines. This has been shown to also result in increased proliferation of the astrocytes of these hiPSC lines.

**NOTE F**: Some hiPSC lines show relatively delayed astrocyte differentiation compared to the majority of the lines (often hiPSC lines being reprogrammed using Sendai vectors). This so-called delayed astrocyte differentiation is defined by hiPSC-like morphology during the first one- or two-weeks during differentiation. This also includes some hiPSC colonies remaining within the astrocyte cultures for the first one or two weeks of differentiation (Supplementary Figure 1E), indicating less efficient astrocyte differentiation for these lines. In order to improve astrocyte differentiation, it is recommended to plate a relative low number of hiPSCs at DIV 0 (100.000 – 150.000 cells per well). In addition, to lose the remaining hiPSC colonies within the astrocyte cultures, it is recommended to split the astrocyte cultures without RevitaCell^TM^.

### Neuronal progenitor cells differentiation

The protocol for the generation of NPCs from hiPSCs was based on a previously published protocol described by Shi et al. (2012)^37^. Briefly, hiPSC colonies were dissociated into single cells using Accutase (Sigma; A6964) and 200.000-300.000 cells (depending on the hiPSC-line) were plated in a Matrigel (Corning; 734-1101) coated 12-well plate (Corning; 353043) and cultured in E8 Flex medium supplemented with RevitaCell™. At 95% cell confluency, which was achieved four days after plating, medium was switched to neural induction medium (NIM). NIM contains 1:1 mixture of Medium A and Medium B medium (referred to as neural maintenance medium (NMM)) supplemented with 1 μM dorsomorphin (Sigma Aldrich; P5499) and 10 μM SB431542 (Stemcell Technologies; 72234) to inhibit SMAD signaling. Medium A medium consists of DMEM/F12, 1x N2 (Gibco; 17502001), 5 μg/mL Insulin (Gibco; 13105002), 2mM GlutaMAX (Gibco; 35050061), 100 μM non-essential amino acids (Thermo Fisher Scientific; 11140-035), 100 μM β-mercaptoethanol (Thermo Fisher Scientific; 21985-023) and Primocin. Medium B is made of neurobasal medium (Gibco; 21103049), 1x B-27 (Gibco; 17504044), 2 mM GlutaMAX and Primocin. For the first ten to twelve days medium was changed every day and the cells were closely monitored for neuroepithelial induction which was based on morphological changes and was expected to start between eight to twelve days after plating. When neuroepithelium was induced (D11), cells were detached from the Matrigel coated plate using ReLeSR™ and transferred onto poly-L-ornithine (PLO; Sigma; P3655)/Biolaminin coated 12-well plates and cultured in NIM supplemented with RevitaCell™. After 24 hours, NIM was changed one more time, which is the last day of incubation with dual SMAD inhibitors (D12). Then medium was switched to NMM supplemented with 10 ng/mL EGF (Peprotech; AF-100-15) and 10 ng/mL FGF (R&D systems; 235-F4-025) and was refreshed every three days. At confluency, after approximately one week of culture, the cells were detached with ReLeSR™ and split onto PLO/Biolaminin coated 6-well plates (P0 of the NPCs). After overnight incubation, the medium was changed to NMM supplemented with 20 ng/mL EGF and 20 ng/mL FGF. From this point on, the cells were passaged at confluency, which was usually four days after passage, in a 1:2 or 1:3 ratio on PLO/Biolaminin pre-coated wells of a 6-well plate in NMM medium supplemented with 20 ng/mL EGF and 20 ng/mL FGF and RevitaCell™. An additional overview of all materials required for the NPC differentiation is provided in Supplementary Table S3. Accordingly, NPC-astrocytes were generated according to the hiPSC-astrocyte protocol.

### Rat astrocytes

The rat astrocytes used in this study were obtained from embryonic E18 rat brains, as described previously^12^. The pregnant rat (Crl:WI[Han], Charles River) underwent anesthesia with isoflurane and was euthanized by cervical dislocation, accordingly the pups were collected and decapitated. The astrocytes were isolated from the brains and pooled. Rats were housed individually since they were euthanized immediately upon arrival at the animal facility. All animal procedures were conducted in accordance with the Animal Care Committee of the Radboudumc, The Netherlands, and adhered to the guidelines of the Dutch Council for Animal Care and the European Communities Council Directive 2010/63/EU (Ethics approval for project W P 2 0 17-0 0 4 8 - 0 0 3).

### Co-culture of iNeurons and hiPSC-derived astrocytes

hiPSCs were differentiated towards excitatory neurons using overexpression of *NGN2* as described previously with minor adjustments^12,38^. Briefly, prior to differentiation, hiPSCs were infected with lentiviral constructs encoding Ngn2 combined with rtTA. In order to obtain a pure population of doxycycline-inducible excitatory neurons, hiPSCs were selected using increasing concentrations of G418 (100 μg/mL - 250 μg/ml; Sigma-Aldrich; #G8168) for the rtTA-construct and puromycin (1 μg/mL - 2 μg/ml; Sigma; P9620) for the Ngn2-construct.

Forty-eight-CytoView (Axion Biosystems) MEA plates or nitric-acid treated glass coverslips (Epredia; 631-0713) were pre-coated with 50 μg/mL PLO diluted in borate buffer for 3 h followed by overnight incubation with 20 µg/mL Biolaminin in 1x dPBS^++^ at 4 °C. To start neuronal differentiation (DIV 0), hiPSC colonies were dissociated into single cells using TrypLE and plated onto the pre-coated MEA plates or coverslips in E8 basal medium (Gibco; A1517001) supplemented with RevitaCell™, doxycycline (4 μg/ml; Sigma; D5207) and Primocin. The next day, medium was refreshed to DMEM/F12 supplemented with N2, 10 ng/mL NT3 (Peprotech; 450-03), 10 ng/mL BDNF (Peprotech; 450-02), MEM non-essential amino acid solution (NEAA; Sigma-Aldrich; M7145), 4 µg/µl doxycycline, and Primocin. Three days after plating (DIV 3) all medium was refreshed to neurobasal medium supplemented with B-27, GlutaMAX, Primocin, NT3, BDNF, and 4 µg/µL doxycycline. In addition, Cytosine β-D-arabinofuranoside (Ara-C; (1 µM; Sigma-Aldrich; C1768) was added to remove proliferating hiPSCs from the culture.

At DIV 6 of the neuronal differentiation, hiPSC-derived astrocytes were added to the neurons. For this the same procedure as described for splitting is applied. Briefly, the hiPSC-derived astrocytes were dissociated using TrypLE and washed by spinning down in DMEM/F12. The pellet was resuspended in neurobasal medium supplemented with B-27, GlutaMAX, Primocin, NT3, BDNF, doxycycline (1 µg/ml; NOTE: lower concentration), RevitaCell™ and 2.5% AM FCS (ScienCell®; 0010). Then medium of the neurons was refreshed with neurobasal medium supplemented with B-27, GlutaMAX, Primocin, NT3, BDNF, doxycycline (1 µg/ml), RevitaCell™ and 2.5% AM FCS. Accordingly, the hiPSC-derived astrocytes were added to the neurons in a neuron:astrocyte ratio 2:1, e.g., 6.750 astrocytes to 13.500 neurons plated on a 48-well CytoView MEA plate. Starting at DIV 8, half of the medium was changed three times a week. To reduce stress for both the hiPSC-derived neurons as well as the hiPSC-astrocytes, doxycycline was withdrawn from the medium starting at DIV 10. The co-cultures were kept at 37 °C/5% CO_2_ through the whole differentiation process. An additional overview of all materials required for the co-culture is provided in Supplementary Table S4.

**NOTE**: It is known that the astrocytes derived from some hiPSC-lines are more sensitive to dissociation than others, leading to an increased cell death upon splitting. Therefore, it is recommended to optimize the number of cells to be used for the co-culture for each hiPSC astrocyte culture.

### Immunocytochemistry

Cells were fixed with 4% paraformaldehyde (Sigma Aldrich; 441244)/4% sucrose (Sigma; S7903) for 15 min, washed three times with PBS (1x; Sigma; P5493) and blocked for 1 h with blocking buffer (BB; 5% normal donkey serum (Jackson Immuno Research; 017-000-121), 5% normal horse serum (Gibco; 26050070), 5% normal goat serum (Invitrogen; 10189722), 1% BSA (Sigma-Aldrich; A-6003), 0.1% D-lysine (Sigma-Aldrich; L802), 1% glycine (Sigma-Aldrich; L802) and 0.4% Triton X-100 (Sigma-Aldrich; T8787) in PBS) at room temperature (RT). Primary and secondary antibodies were diluted in BB and incubated overnight at 4 °C or for 1 h at RT, respectively. To stain the nuclei, cells were incubated with Hoechst (Thermo Scientific; 62249) diluted in PBS for 10 min at RT and accordingly mounted using DAKO fluorescent mounting medium (DAKO; S3023). Zeiss Axio Imager Z1 was used to image the samples. The following antibodies were used: Rabbit anti-Vimentin (1:300; Abcam; ab92547), Mouse anti-Tuj1 (1:300; BioLegend; 801201), Mouse anti-Nestin (1:200; Invitrogen; MA1110), Rabbit anti-EAAT1 (1:100; Abcam; ab416), Rabbit anti-GFAP (1:300; Sigma-Aldrich; AB5804), Mouse anti-CD44 (1:300; Invitrogen; MA5-13890), Mouse anti-ALDH1L1 (1:300; Novus Biologicals; NBP2-50045), Rabbit anti-GLUD1 (1:300; Invitrogen; PA5-28301), Mouse anti-AQP4 (1:300; Biorbyt; ORB323095), Rabbit anti-PDLIM7 (1:300; Novus Biologicals; NBP2-58734), Guinea pig anti-MAP2 (1:1000; Synaptic Systems; 188 004), Mouse anti-Synapsin I (1:1000; Synaptic Systems; 106 001), Goat anti-Rabbit Alexa Fluor 568 (1:1000; Molecular probes; A11011), Goat anti-Mouse Alexa Fluor 488 (1:1000; Invitrogen; A11029), Goat anti-Guinea Pig Alexa Fluor 488 (1:1000; Invitrogen; A11073), Goat anti-Guinea Pig Alexa Fluor 647 (1:1000; Molecular probes; A21450), Goat anti-Mouse Alexa Fluor 647 (1:1000; Molecular probes; A21237), Streptavidin, Alexa Fluor™ 568 conjugate (1:1000; Invitrogen; S11226).

### Flowcytometry analysis

At DIV 35, astrocytes were dissociated into a single-cell suspension using TrypLE. The cells were dissolved in Viability Dye eFluor 780 (Invitrogen; 65-0865-14,) and incubated at 4 °C for 30 min. Subsequently, cells were washed in flow buffer by centrifugation at 300xg for 5 min (1% BSA in PBS) and then fixed for 15 min at RT with 2% paraformaldehyde followed by another wash. To prevent a specific binding to Fc-receptors, the cells were blocked with 2.5% Human TruStain FcX blocking solution (BioLegend; 422302) for 10 min. Subsequently, the cells were incubated with anti-CD44 Brilliant Violet 421 (Biolegend; 338809) and CD49f PE/Cyanine7 (Biolegend; 313621) for 30 min at 4 °C. Afterwards, the cells were washed two times and finally dissolved in 200 µl flow buffer for flow cytometry measurement. Gating of side scatter (SS) area versus forward scatter (FS) area was used to define a cell population based on size and granularity. To remove duplicates, gating between FS height and FS Area was used. Further, dead cells were negatively gated by viability dye intensity (Invitrogen; 17324361. This dye reacts with free amines and gives an intense fluorescent signal when the membrane is ruptured. In live cells the reactivity of the dye is restricted to cell membrane, which results in distinctively different fluorescence. An unstained sample was used to determine the auto-fluorescence levels of cells in the channels corresponding to the antibodies used. This was used to set the threshold for the CD49f- and CD44-positive populations. Analysis was performed using Kaluza C Analysis software (Beckman Coulter).

### RT-qPCR

Total RNA was isolated from the cells with NucleoSpin RNA Mini Kit (Macherey-Nagel; MN 740955.250), according to the manufacturer’s instructions, with the following adjustments: no use of β-mercaptoethanol, and 30 min instead of 15 min incubation with rDNase. Subsequently, RNA was reverse-transcribed into cDNA using iScript cDNA Synthesis Kit (Bio-Rad; 1708891). GoTaq quantitative PCR (qPCR) Master Mix (Promega; A6002) was used to conduct the qPCR. The following program was used in the 7500 Fast Real Time PCR System apparatus (Applied Biosystems): a denaturation at 95 °C for 2 min, 40 cycles of 30 sec 95 °C and 30 sec 60 °C followed by a melting curve stage of 15 sec 95 °C, 30 sec 60 °C and 15 sec 95 °C. Samples were always prepared in triplicate. Outliers were determined if a value differed more than 0.5 Ct from the other two values in the technical triplicate. The relative mRNA expression was calculated using the 2−ΔΔCt method with normalization against the housekeeping genes *GUSB, ERCC6* and *PPIA*. Primers are listed in Supplementary table S5.

### RNA sequencing

#### RNA-seq library preparation

Five different hiPSC-derived astrocytes cultures (hiPSC-line 1, hiPSC-line 11, hiPSC-line 4, hiPSC-line 6 and hiPSC-line 7) and three different NPC-derived astrocyte cultures (hiPSC-line 1, hiPSC-line 2 and hiPSC-line 15) were prepared for RNA sequencing analysis. Biological triplicates of all astrocyte cultures were seeded onto 6-well plates in AM. Upon confluency, the astrocytes were washed twice with ice-cold PBS and collected using DNA/RNA shield (Zymo 20 Research; #ZY-R1200-125). Accordingly, RNA was isolated using Quick-RNA Microprep kit (Zymo 20 Research, #ZY-R1051), according to manufacturer’s instructions. The quality of RNA was assessed using Agilent’s Tapestation system and the RNA Integrity Number (RIN) values ranged between 7.3 and 9.6. Subsequently, cDNA libraries were prepared using the NEBNext Ultra II Directional RNA Library Prep Kit, and paired-end reads were sequenced on an Illumina NovaSeq 6000 platform at GenomeScan B.V. Leiden.

#### RNA-seq data processing

We used Fastp to eliminate PolyG artifacts and clip adapters, including a list of adapter sequences currently used by Illumina^39^. Subsequently, we mapped the reads to the GRCh38 human reference genome using HISAT2. As the library was reversely stranded, we set --rna-strandness = RF. The resulting SAM files were sorted and indexed using SAMtools^40^. Next, we performed UMI deduplication with UMI-tools^41^, and feature counting was done with Subread’s *featureCounts*^42^.

Paired-end RNA-seq data of *post-mortem* human astrocytes were downloaded from GSE73721 GEO repository^43^. Fastp was used to trim the reads, while HISAT2 was employed to align the reads to the GRCh38. SAM files were sorted and indexed with SAMtools, and *featureCounts* for feature counting.

#### RNA-seq analysis

Raw count matrices were loaded in R version 4.2.1. When Ensemble IDs mapped to the same gene symbols, we considered only the IDs with the highest expression per sample. We filtered out genes with low expression by retaining those with expression values exceeding one transcript per million reads in at least six samples. Subsequently, counts were normalized using *voom* function from the limma R package^44^. Genes were then ranked based on their variance, and Principal component analysis (PCA) was performed on scaled expression data of the 2500 most variable genes, using *prcomp* function. Given that the first principal component predominantly explained the variability between human *post-mortem* and *in-vitro* samples, the Euclidean distance between each pair of samples was calculated based on the values of the principal components, excluding the first component. These distances were then used for hierarchical clustering with *pheatmap* function of pheatmap R package. To create heatmaps illustrating the expression patterns of astrocyte marker genes across the samples, normalized counts were scaled between genes. All plots were generated using custom code built on ggplot2 v.3.4.0 functions in R.

### Glutamate uptake analysis

To prepare samples for glutamate uptake analysis, 20.000 hiPSC-astrocytes (DIV 34) or rat astrocytes per well were seeded onto 24-well plates (Corning; 353047) and cultured in AM overnight at 37 °C/5% CO_2_. Three hours prior to harvest, AM supplemented with 100 µM glutamic acid (Sigma; G1251) and Primocin was refreshed. Upon incubation the medium was collected and stored at −20 °C until further processed. To measure uptake of glutamic acid by the hiPSC-derived astrocytes, samples were diluted 50 times and measured using the Amplex Red Glutamic Acid/Glutamate Oxidase Assay Kit (Thermo Fisher Scientific; A12221,) according to manufacturer’s instructions. Fluorescent signal was measured at 540 nm excitation and 590 nm emission using a microplate reader (Tecan’s Spark® 20M multimode reader). The percentage of glutamic acid taken up by the astrocytes relative to the negative control was calculated.

### Biocytin filling using patch-clamping

Coverslips with hiPSC-astrocyte monocultures were placed in a recording chamber on the stage of an Olympus BX51WI upright microscope (Olympus Life Science), equipped with infrared differential interference contrast optics, an Olympus LUMPlanFL N 60x water-immersion objective (Olympus Life Science), and a kappa MXC 200 camera system (Kappa optronics GmbH) for visualization. Through the recording chamber there was a continuous flow of carbongenated (95% O_2_/5% CO_2_) artificial cerebrospinal fluid (aCSF) (124 mM NaCl (Sigma; S5886), 1.25 mM NaH_2_PO_4_ (Sigma; S0751), 3 mM KCl (Sigma; P9333), 26 mM NaHCO_3_ (Sigma; S5761), 11 mM Glucose (Sigma; G5767), 2 mM CaCl_2_ (Merck; 1023780500), 1 mM MgCl_2_ (Merck; M2670)) at 32°C. Patch pipettes (ID 0.86 mm, OD1.05 mm, resistance 5–8 MΩ) were pulled from borosilicate glass with filament and fire-polished ends (Science Products GmbH) using the Narishige PC-10 micropipette puller. Pipettes were filled with a potassium-based solution with 0.5% biocytin (Sigma-Aldrich; B4261) containing: 130 mM K-Gluconate, 5 mM KCl, 10 mM HEPES, 2.5 mM MgCl_2_, 4 mM Na_2_-ATP, 0.4 mM Na_2_-GTP, 10 mM Na-phosphocreatine and 0.6 mM EGTA (with pH adjusted to 7.2 and osmolarity to 290 mOsmol). Passive properties were continuously monitored using a Digidata 1140A digitizer and a Multiclamp 700B amplifier (Molecular Devices). After reaching whole-cell configuration, astrocytes were kept at a holding potential of −60 mV for 15 min to allow the biocytin to diffuse out of the pipette into the attached astrocytes. After 15 min the pipette was slowly retracted to reseal the plasma membrane. Coverslips were left in the recording chamber for another 15 min to wash off excess biocytin.

### Calcium imaging

To prepare the cells for calcium imaging analysis, hiPSC-derived astrocytes, grown on glass coverslips, were incubated with Fluo-8-AM calcium binding dye (Abcam; ab142773) diluted in HBSS buffer (Sigma; H6648) supplemented with 20 mM HEPES (Sigma-Aldrich; H4034) for 30 min at 37 °C. To remove excessive Fluo-8-AM, the cells were washed three times with HBSS buffer and subsequently allowed to recover in AM without FBS for 15 min. To image the hiPSC-derived astrocytes, the OLYMPUS BX51WI Upright Microscope using a 20x magnification was used combined with the Hamamatsu camera (ORCA Flash2.8). During recordings, the cells were illuminated at 471nm by LED light (KSL470, Rapp OptoElectronic). The cells were perfused continuously with oxygenated aCSF (119 mM NaCl, 26.2 mM NaHCO3, 2.5 mM KCl, 1 mM NaH2PO4, 21 mM glucose, 2 mM CaCl2, 2.3 mM MgCl2) and measured in aCSF supplemented with or without 50 µM glutamic acid. Fluorescent signals were recorded for 10 min at 250 msec intervals (4Hz). During evoked recordings, the cells were recorded for 2 min to obtain the baseline measurements. Accordingly, while recording, the original aCSF was replaced by aCSF supplemented 50 µM glutamic acid, which was slowly perfused towards the cells.

Data were analyzed using a home-made script in MATLAB (The Math Works, Inc. MATLAB. Version 2020b). Regions of interest (ROIs) were selected manually per measurement to represent single hiPSC-derived astrocytes. Subsequently, the mean signal of each ROI was plotted per frame. The calcium response signal was calculated as the change in fluorescent signal of the ROI according to the formula ΔF/F = (F-F0)/F0. F represents the fluorescent signal for each ROI over time, while F0 represents the baseline fluorescence. To correct for the decay of the baseline intensity due to bleaching, exponential fitting was used. To represent the mean intensity for each ROI, the area under the curve was calculated.

### Olink® proteomics

To assess the astrocyte secretome, DIV 35 hiPSC-line 1 astrocytes were sampled for Olink® proteomics using the Target 96 inflammation panel. The Olink® assay is based on Proximity Extension Assay (PEA) technology. In short, 96 different oligonucleotide antibody-pairs are designed which can bind to the proteins in the sample. These antibody-pairs contain unique DNA sequences, which allow for hybridization to each other upon specific binding to the protein. Subsequent proximity extension will create 96 unique DNA reporter sequences which are amplified by real-time PCR.

Astrocytes were seeded onto 6-well plates several days prior to sample collection to allow the cells to grow confluent. The day before sample collection, the medium was refreshed. At DIV 35 medium of the astrocytes was collected and stored at −20 °C until further processed. In addition, as baseline control fresh medium was collected and stored. The Olink® assay was performed according to the manufacturer’s instructions. Several controls were included. The inter-plate Control (IPC) is a pool of 92 antibodies, each with one pair of unique DNA-tags positioned in fixed proximity. In addition, a negative control was included consisting of running buffer, as well as a sample control consisting of pooled samples to assess potential variation between runs. The data were first normalized for the IPC. Afterwards the fold change for each protein was calculated relative to the baseline medium sample.

### Synapsin quantification

To quantify the number of Synapsin puncta, hiPSC-astrocyte and iNeuron co-cultures were stained for MAP2, Vimentin and Synapsin. Ten images per coverslip were made using Zeiss Axio Imager Z1. Accordingly, the number of Synapsin puncta per X µm dendrite were quantified using FIJI. Synapsin puncta for two neurons (one or two dendrites) per image were calculated. Accordingly, the number of Synapsin puncta per 10 µm dendrite was calculated.

### MEA recordings and analysis

To record spontaneous network activity of human co-cultures using MEAs, hiPSC-derived neurons and hiPSC-astrocytes were plated onto the CytoView 48-well MEA plates (Axion Biosystems) as described previously. At least four wells per culture condition were included for each experiment. In brief, each well contains 16 electrodes with a diameter of 50 µm arranged in a 4 x 4 grid spaced 350 µm apart. Recordings were performed weekly, starting at DIV 21 until DIV 51 of the differentiation. Spontaneous network activity was recorded for 5 min, after a 5 min period of acclimatization, in a recording chamber which was maintained at 37 °C/5% CO2. Spike threshold at which spikes were detected was set at +-6 STD.

Analysis of the MEA recordings was performed using the Envelope burst parameter of the Axion Biosystems Neural Metric Tool. Threshold factor was set at 3.0, minimum inter burst (ms) at 100ms, and both the minimum number of active electrodes and burst inclusion was set at 50%. From the analysis the following parameters were extracted: mean firing rate, the number of bursts, the number of network bursts (NBs) and the NB duration.

### Patch clamp

Single cell recordings in whole cell patch clamp configuration were performed at DIV35 and as previously described^38,45^. Coverslips were placed in a recording chamber on the stage of an Olympus BX51WI upright microscope (Olympus Life Science), equipped with infrared differential interference contrast optics, an Olympus LUMPlanFL N 60x water-immersion objective (Olympus Life Science), and a kappa MXC 200 camera system (Kappa optronics GmbH) for visualization. The recording chamber was continuously perfused with carbongenated (95% O_2_/5% CO_2_) aCSF (124 mM NaCl, 1.25 mM NaH_2_PO_4_, 3 mM KCl, 26 mM NaHCO_3_, 11 mM Glucose, 2 mM CaCl_2_, 1 mM MgCl_2_) at 32 °C. Patch pipettes (ID 0.86 mm, OD1.05 mm, resistance 5–8 MΩ) were pulled from borosilicate glass with filament and fire-polished ends (Science Products GmbH) using the Narishige PC-10 micropipette puller. For recordings of intrinsic properties in current clamp mode and spontaneous excitatory postsynaptic currents (sEPSCs) in voltage clamp mode, pipettes were filled with a potassium-based solution containing: 130 mM K-Gluconate, 5 mM KCl, 10 HEPES, 2.5 mM MgCl_2_, 4 mM Na_2_-ATP, 0.4 mM Na_3_-GTP, 10 mM Na-phosphocreatine, 0.6 mM EGTA (with pH adjusted to 7.2 and osmolarity to 290 mOsmol). All recordings were acquired using a Digidata 1140A digitizer and a Multiclamp 700B amplifier (Molecular Devices), with a sampling rate set at 20 kHz and a lowpass 1kHz filter during recording. Recordings were not corrected for liquid junction potential (±17 mV), which was calculated according to the stationary Nernst–Planck equation^46^ using LJPcalc software (https://swharden.com/LJPcalc). Recordings were not analyzed if series resistance was above 25 MΩ or when the recording reached below a 10:1 ratio of membrane to series resistance. Resting membrane potential (Vrmp) was determined directly after reaching whole-cell configuration. Further analysis of active and passive membrane properties was conducted at a holding potential of −60 mV. sEPSCs were measured by 10 min continuous recording in aCSF at a holding potential of −60 mV. Amplitude and frequency of individual events within sEPSC recordings were not analyzed using Mini Analysis.

### Statistical analysis

Statistical analysis was performed using Graphpad Prism (version 10 for Windows, GraphPad Software). For comparisons between two conditions at one time point, an unpaired t-test was performed. All values are reported as mean ± standard deviation (SD). Results with P values lower than 0.05 were considered as significantly different (*), P < 0.01 (**), P < 0.001 (***), P < 0.0001 (****).

## Results

### Direct differentiation from hiPSCs towards morphologically mature astrocytes

In this study thirty hiPSC-lines (Supplementary Table S1) were differentiated towards astrocytes in different batches (Supplementary Figure 2 and 3). For a subset of these hiPSC-astrocyte cultures, we performed in-depth characterization to validate the protocol and to confirm astrocyte identity. Astrocyte differentiation was initiated by plating the hiPSCs as single cells on human Biolaminin®-coated plates and culturing them in commercial AM (ScienCell®) supplemented with RevitaCell^TM^ (Figure 1A, Methods). After 48 h of plating, we already observed the first changes in morphology as the cells exhibited a more flat and less condensed appearance compared to hiPSCs. Over the course of the first week of differentiation, these morphological alterations persisted, with cells gradually increasing in size, flattening and demonstrating reduced proliferation rates relative to hiPSCs. By the end of the initial three weeks, the astrocytes formed a predominantly homogenous culture. However, the astrocytes changed into a more heterogeneous culture, characterized by differences in astrocyte size and morphology, starting after approximately three weeks (depending on the hiPSC-line) (Figure 1A). To further characterize morphological maturation and astrocytic identity over time, the hiPSC-astrocytes were immunostained for vimentin and Tuj1 at multiple time points (Figure 1B). While vimentin is generally considered a mature astrocyte marker, Tuj1 is associated with neuronal identity, albeit it has been reported to be expressed by early astrocytes as well^47^. We found that the expression of vimentin increased, while the expression of Tuj1 decreased during the differentiation process. We also found increased morphological heterogeneity over time, which was evidenced by differential expression of vimentin and Tuj1 and by increased astrocyte size as well as morphological complexity. This was most strikingly observed by the increasing number of astrocytic processes during maturation.

**Figure 1.**
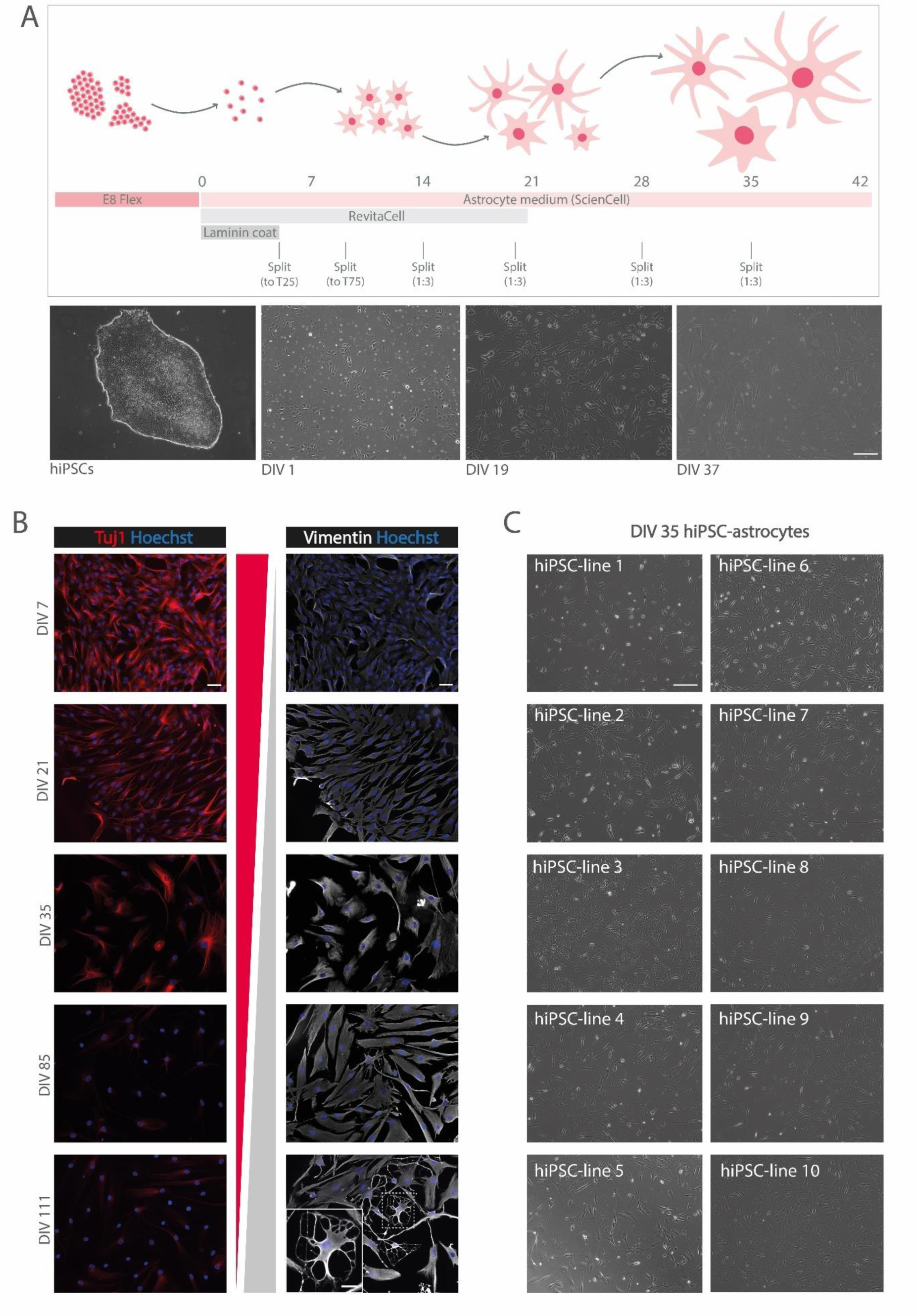
Direct differentiation from hiPSCs towards astrocytes results in morphological mature astrocytes. **(A)** Schematic representation of the protocol to directly differentiate hiPSCs towards astrocytes, including brightfield images over time during differentiation. All pictures were taken at the same magnification (scale bar = 100 µm). **(B)** Immunostaining of vimentin (white) and Tuj1 (red) of hiPSC-astrocytes (hiPSC-line 4) at DIV 7, DIV 21, DIV 35, DIV 85 and DIV 111, showing decreased and increased protein level of Tuj1 and vimentin over time, respectively. All pictures were taken at the same magnification (scale bar = 50 µm or 25 µm in zoom image). **(C)** Brightfield images of ten different DIV 35 hiPSC-astrocyte cultures of one batch. All pictures were taken at the same magnification (scale bar = 100 µm).

Among the thirty differentiated hiPSC lines (Supplementary Table S1), we found line-to-line variability, defined as differences in morphology, size and proliferation rate, across different batches (Supplementary Figure 1-3, Figure 1C). Importantly, the astrocyte differentiation trajectory for each hiPSC-line was highly consistent across different batches, as shown by similar morphological characteristics across different astrocyte batches using brightfield and immunofluorescent imaging (Supplementary Figure 4). These results demonstrate minimal batch-to-batch variability, confirming the robustness of this astrocyte differentiation protocol. We found no direct correlation between morphological differences or proliferation rate and the quality or functionality of the hiPSC-astrocytes. Notably, we observed a delayed astrocyte differentiation (defined by hiPSC-like morphology during the first one- or two-weeks during differentiation) in hiPSC lines reprogrammed using Sendai vectors compared to those reprogrammed using lentiviral or episomal vectors. However, additional experiments featuring systematic side-by-side comparisons would be necessary to further validate this observation.

### Expression of functional astrocyte markers confirm maturation of hiPSC-derived astrocytes

To assess the purity and cell-lineage specificity of the hiPSC-derived astrocyte cultures, we performed flow cytometry on DIV 35 hiPSC-astrocytes (Figure 2A). Flow cytometry analysis showed differences in astrocyte size accounting for morphological astrocyte heterogeneity as previously described^48,49^. In addition, the astrocytes were highly viable (92%) and showed expression of CD49f, indicating successful differentiation towards the astrocyte lineage^50^. We observed a subpopulation that was negative for CD44 (29%) which further demonstrates cellular heterogeneity. Seventy percent of the hiPSC-astrocytes co-expressed CD44 and CD49f at DIV 35.

**Figure 2.**
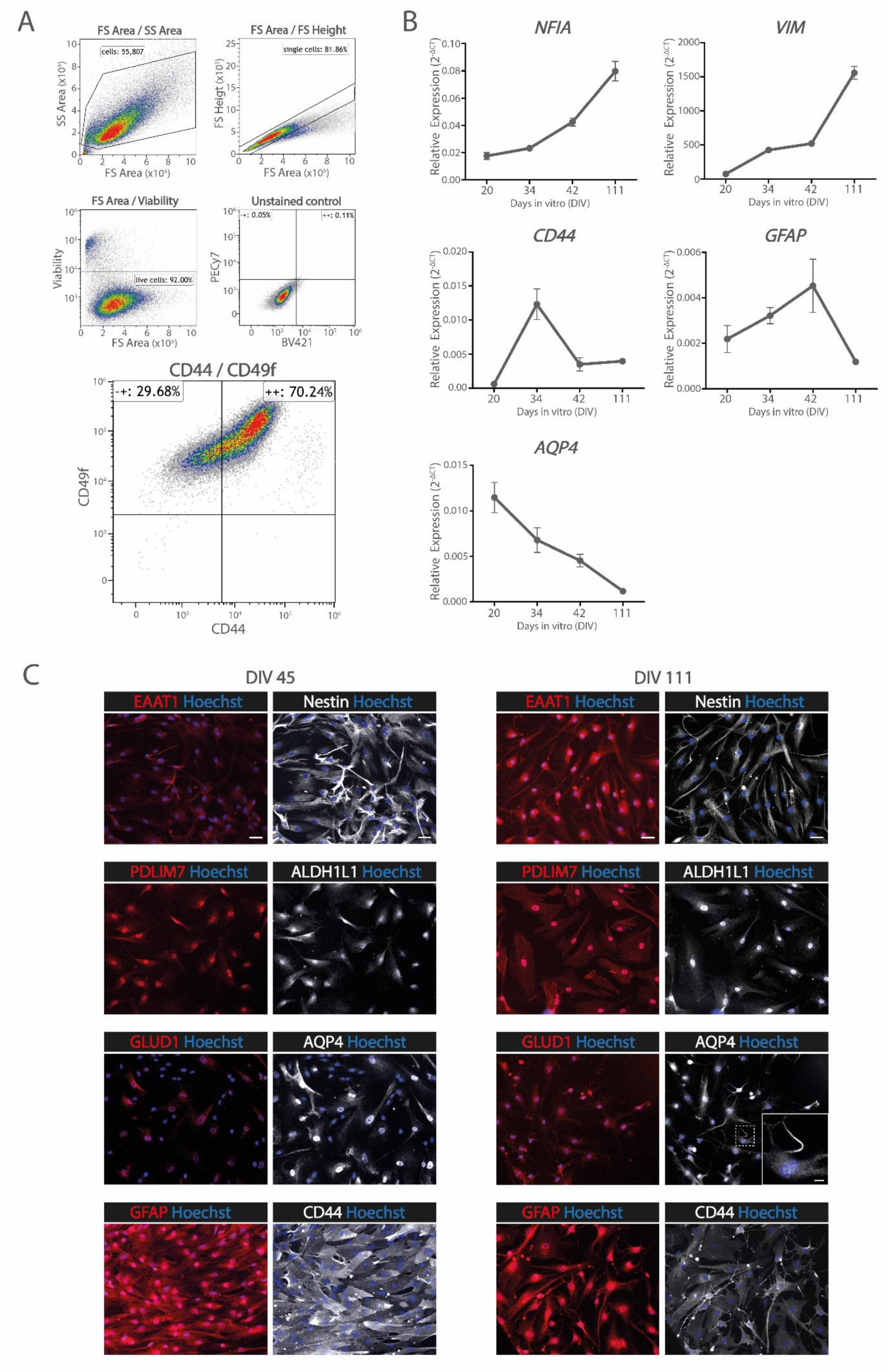
Molecular characterization of hiPSC-astrocytes confirms astrocyte maturation and identity. **(A)** Flow Cytometry data, including viability and unstained control, showing co-expression of CD44 and CD49f by 70% of the hiPSC-astrocyte population. **(B)** Relative expression of *NFIA, VIM, CD44, GFAP* and *AQP4* by qPCR in DIV 20, DIV34, DIV 42 and DIV 111 hiPSC-astrocytes (hiPSC-line 1). Expression of genes was normalized for the average expression of *GUSB, ERCC1* and *PPIA*. **(C)** Protein level of the functional astrocyte markers EAAT1 (red), Nestin (white), PDLIM7 (red), ALDH1L1 (white), GLUD1 (red), AQP4 (white), GFAP (red) and CD44 (white) in DIV 45 and DIV 111 hiPSC-astrocytes. All pictures were taken at the same magnification (scale bar = 50 µm or 10 µm in the zoomed image).

To track the development of hiPSC-astrocytes over time, we performed qPCR for *NFIA, VIM, CD44, GFAP* and *AQP4* at various time points: DIV 20, DIV 34, DIV 42 and DIV 111 (Figure 2B). As expected, we observed increased expression of *NFIA* and *VIM* over time. Expression of *CD44* peaked at DIV 34 after which it decreased. *GFAP* expression increased for the first three timepoints but was decreased at DIV 111. Interestingly, we find that *AQP4* expression decreased over time. This could be explained by the fact that at early timepoints the AQP4 protein level is high as it is localised through the whole cell, while during maturation and therefore at later timepoints the AQP4 protein level is decreased as it is localised and limited to the astrocytic end feet^51–53^, as observed by immunofluorescence (Supplementary Figure 5).

We further confirmed a high protein level of astrocyte specific markers by DIV 45 and DIV 111 hiPSC-astrocytes using immunofluorescence (Figure 2C). We observed high protein levels of Nestin and ALDH1L1 at both time points. The protein level of EAAT1, GLUD1 and PDLIM7 increased over time. PDLIM7 also specifically localised to the actin filaments at DIV 111. Similar as we observed by qPCR, the protein level of GFAP and CD44 was higher at DIV 45 compared to DIV 111. As mentioned before, AQP4 was present throughout the whole cell body in most DIV 45 astrocytes, while at DIV 111 AQP4 was mostly limited to the astrocytic end feet (Supplementary Figure 5).

### RNA sequencing confirms astrocytic identify of hiPSC-astrocytes (and NPC-astrocytes)

As far as we know, all *in vitro* hiPSC-astrocyte differentiation protocols in literature first differentiate the hiPSCs towards NPCs or progenitor-like cells e.g., by dual SMAD inhibition, which are differentiated towards astrocytes in a second step. Our protocol did not include such an intermediate differentiation step, instead we directly differentiated hiPSCs to astrocytes. To asses the molecular similarity between hiPSC-derived astrocytes, NPC-derived astrocytes, and primary human astrocytes, as well as other brain cell types, we performed bulk RNA sequencing for biological triplicates of five different hiPSC-astrocyte cultures and three different NPC-astrocyte cultures^20,37^. Additionally, we included previously published RNA sequencing data of post-mortem fetal and adult astrocytes, along with other CNS cell types^43^. Principal component (PC) analysis showed that the hiPSC-astrocytes and NPC-astrocytes clustered closely together (Figure 3A-B), demonstrating minimal differences between direct differentiation from hiPSCs towards astrocytes and astrocyte differentiation including an intermediate NPC stage. While the first PC separated the samples based on their origin (*in-vitro* or *post-mortem*) (Figure 3A), the second and third PCs segregated the samples regarding their cell type. So all astrocytes, irrespective of their origin, clustered more closely together compared to the other brain cell types (Figure 3B). Hierarchical clustering of the samples based on PCs-derived distances further supported this observation, grouping *post-mortem* and *in-vitro* astrocytes closer together in comparison to neurons, oligodendrocytes and myeloid cells (Figure 3C). We observed that astrocyte specific genes such as *VIM, GJA1, CD44, PDLIM7, TNC, APOE, GLUD1, GLUL* and *NFIX* are highly expressed by both the hiPSC-astrocytes and NPC-astrocytes, while neuronal genes (*DCX, NGN2* and *RBFOX3*) and oligodendrocyte genes (*SOX10* and *MBP*) are lowly expressed or not detected (Figure 3D). The expression of hiPSC genes (*OCT4, SSEA4*) was not detected (data not shown). However, other astrocyte specific genes, such as *GFAP* and *S100B* and glutamate transporter genes such as *SLC4A4* and *SLC1A2* were expressed at relatively low levels by the hiPSC-astrocyte monocultures.

**Figure 3.**
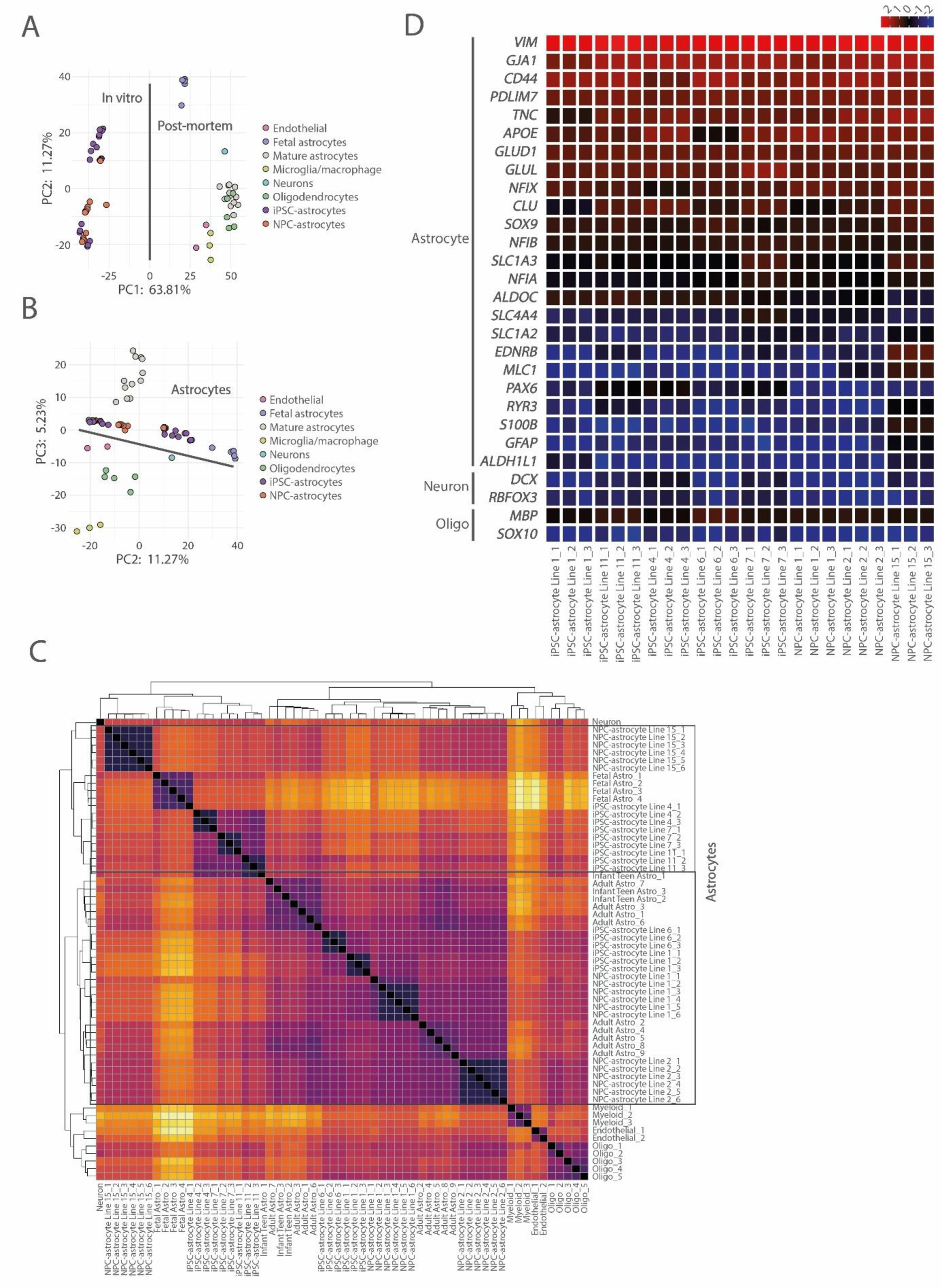
RNA sequencing analysis of hiPSC-astrocytes and NPC-astrocytes compared to post-mortem astrocyte samples. Bulk RNA sequencing of five different hiPSC-astrocyte cultures and three different NPC-astrocyte cultures compared to post-mortem astrocyte gene expression datasets. Principle component (PC) analysis was performed of which **(A)** PC1 and PC2 showed separation based on origin and **(B)** PC2 and PC3 showed separation based on cell type identity, showing clustering of hiPSC-astrocytes and NPC-astrocytes with post-mortem astrocyte samples. **(C)** Hierarchical clustering based on PCs-derived sample distances. **(D)** Heatmap gene expression by the hiPSC-astrocytes and NPC-astrocytes, indicating high expression of astrocyte genes, and low expression of neuronal and oligodendrocyte genes.

### Functional characterization of hiPSC-astrocytes confirms their ability to take up glutamate and form gap junctions in monoculture

One of the main functions of astrocytes is to take up excessive glutamate from the synaptic cleft. To assess this functionality, we measured the percentage of glutamic acid uptake by DIV 35 monoculture hiPSC-astrocytes three hours after adding glutamic acid to the medium. We assessed ten different hiPSC-astrocyte cultures and one rat astrocyte culture (Supplementary Figure 6). Three hiPSC-astrocyte cultures showed 20-30% uptake of the additional glutamic acid, which is 2-3 times higher compared to the rat astrocytes. Four other hiPSC-astrocyte cultures showed 5-10% uptake, while three hiPSC-astrocyte cultures showed only a minimal uptake of glutamic acid. We continued further functional characterization with hiPSC-line 1 astrocytes as they showed most effective uptake of glutamic acid (Figure 4A).

**Figure 4.**
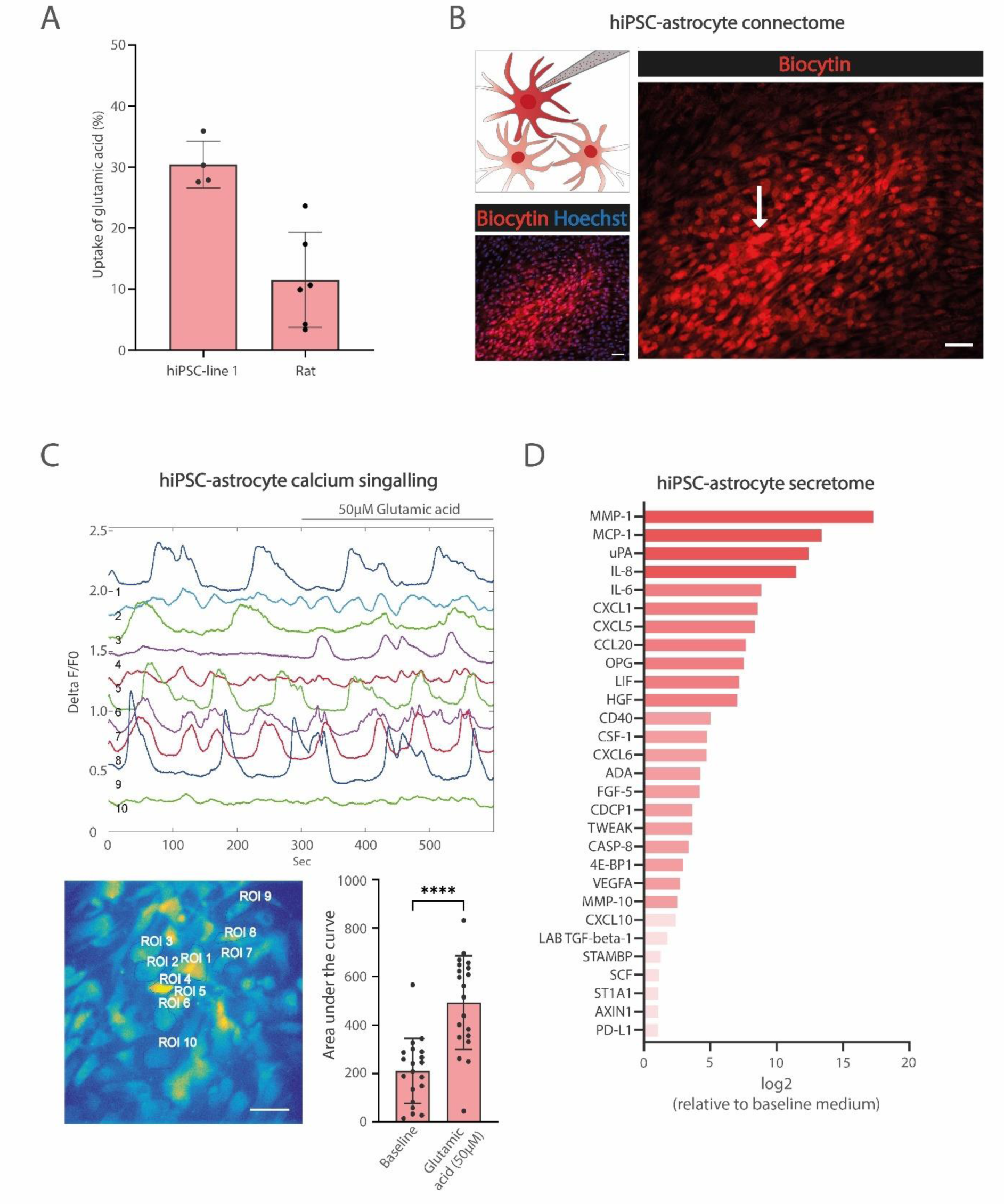
Functional characterization of hiPSC-astrocytes confirms the ability to take up glutamate and communicate. **(A)** The percentage of glutamic acid uptake by monoculture hiPSC-line 1 astrocytes was measured and compared to one rat astrocyte culture. **(B)** Astrocyte connectivity was assessed by injection of Biocytin using single-cell patch-clamping of one hiPSC-astrocyte in monoculture, followed by immunolabeling with a Streptavidin 568 conjugated antibody. **(C)** Astrocytic Ca^2+^ signaling was measured upon incubation with the Fluo-8-AM dye, by 10 min recording of which the first 5 min were baseline followed by a 50 µM glutamic acid stimulus. Astrocytic Ca^2+^ signaling was calculated as area under the curve of the calcium waves. One-way ANOVA showed significantly increased activity upon glutamic acid stimulus. Significance is indicated by **** (P < 0.0001) (scale bar = 50 µm). **(D)** The astrocyte secretome was analyzed using Olink® proteomics. Data were first normalized for the interplate control, and accordingly fold change for each protein was calculated relative to baseline medium. Proteins with >1 fold change are showed.

To facilitate rapid communication among astrocytes, they are interconnected through gap junctions, forming what is known as the astrocytic syncytium or astrocyte connectome. Previous *in vivo* studies have delineated the connectivity of astrocytes in the brain^54,55^. To evaluate the capacity of our hiPSC-astrocytes in monoculture to establish gap junctions and interconnect, we utilized biocytin, a cellular tracer, which was injected into one astrocyte via single cell path-clamping. The wide spreading of the biocytin dye demonstrated the ability of hiPSC-astrocytes to establish connections (Figure 4B).

Astrocytic communication often involves Ca^2+^ signaling. Various stimuli such as ATP or glutamic acid can trigger astrocytic Ca^2+^ signaling, leading to the release of gliotransmitters^56^. To assess the astrocytic Ca^2+^ signaling ability of our hiPSC-astrocytes in monoculture, we employed Fluo-8-AM labeling. We first recorded baseline activity followed by a stimulation with glutamic acid (Figure 4C). At baseline, most astrocytes were active, of which some astrocytes demonstrated synchronized calcium waves. Upon stimulation with glutamic acid, astrocytic Ca^2+^ signaling exhibited a significant increase, confirming their ability to rapidly respond to stimuli.

### hiPSC-astrocytes secrete growth factors involved in neuronal and glial development

The importance of astrocyte-secreted proteins on the development, maturation and function of neurons and synapses is well described in literature^57,58^. These proteins support neuronal function and development, but are also crucial for the development and maturation of astrocytes and other glial cell types. To assess the astrocyte secretome we performed Olink® proteomics on hiPSC-astrocytes in monoculture (Figure 4D). We observed a relative high secretion of proteins affecting neuronal function and development, such as MCP-1, IL-6, HGF, TWEAK and VEGFA^59–63^. In addition, some of the highly secreted proteins such as IL-6 and CSF-1 are known to be important for both microglia or oligodendrocyte development^64,65^. LIF, an important growth factor known to be involved in astrocyte development^66,67^, was also highly secreted by the hiPSC-astrocytes. Interestingly, in some astrocyte differentiation protocols, LIF is described as one of the main drivers of astrocyte differentiation^20,68^.

### Optimization of the protocol to co-culture hiPSC-astrocytes together with iNeurons

Our primary motivation to develop a new astrocyte differentiation protocol was to establish an all-human astrocyte-neuron co-culture, in which astrocytes functionally support neuronal development. We cultured iNeurons according to our previously published protocol^12^ and assessed the ability of the hiPSC-astrocytes to support neuronal development. Our key criteria for evaluating the co-culture protocol included robustness and reproducibility as well as high-quality neurons, defined by fine dendritic outgrowth and no neuronal clustering. Immunofluorescent imaging and MEAs served as our primary measures for assessing co-culture viability and functionality.

Unfortunately, utilizing the standard co-culture protocol^12^ did not result in viable and reproducible co-cultures (Supplementary Figure 7). In order to improve viability of the co-cultures, the following adjustments to the protocol were tested: varying concentrations of doxycycline; different timing of adding FCS as well as different types of FCS; various types of medium during plating of the astrocytes; addition of RevitaCell^TM^ during plating of the astrocytes; withdrawal of AraC. The effect of these adjustments on the viability of both cell types was assessed using immunofluorescent imaging. Upon implementation of the adjustments into the protocol (addition of Revitacell while plating of the astrocytes; lower doxycycline concentration when astrocytes are added; standard FBS replaced by AM FBS to lower the endotoxin concentrations), immunofluorescent imaging confirmed significant improved viability of both the iNeurons and hiPSC-astrocytes in the co-cultures (Supplementary Figure 7).

To investigate whether these iNeurons are also active and form synchronized neuronal networks when co-cultured with hiPSC-astrocytes, we measured spontaneous neuronal activity using MEAs. Synchronized neuronal network activity, which is defined by NBs, are generally considered a characteristic for mature *in vitro* neuronal cultures^69–73^. Therefore, while assessing the co-culture protocol, the number of NBs was considered as the most important parameter. We measured neural activity of the hiPSC-line 1 co-cultures on MEAs including four biological replicates. While the appearance of NB activity was considered positive, the NB frequency was highly variable across the four different replicates (Supplementary Figure 7). We therefore implemented additional adjustments to the protocol (neuron:astrocyte ratio 2:1 instead of 1:1; DIV 42 instead of DIV 35 hiPSC-astrocytes; addition of FBS on DIV 3 instead of DIV 10 to promote astrocyte survival) and assessed these co-cultures on the MEA. Upon the new adjustments only 50% of the biological replicates showed NB activity, which also showed varying NB frequency and NB duration (Supplementary Figure 7). Upon implementation of the final adjustments (adding hiPSC-astrocytes at DIV 6 instead of DIV 3; more details in Methods) all four biological replicates showed a consistent number of NBs and NB duration (Supplementary Figure 7).

### hiPSC-astrocytes support neuronal development, maturation and synapse formation when co-cultured with iNeurons

Upon validating the hiPSC-astrocytes in monoculture, we assessed whether the hiPSC-astrocytes also functionally support neuronal development when co-cultured with iNeurons. Our key criteria for evaluating the co-culture protocol included robustness and reproducibility as well as high-quality neurons, defined by fine dendritic outgrowth and no neuronal clustering. To confirm the reproducibility of the hiPSC-astrocytes within the co-cultures, we tested multiple hiPSC-lines across different batches, and assessed these co-cultures using immunocytochemistry (Figure 5A). The overall viability and quality of both the iNeurons and hiPSC-astrocytes in all co-cultures was high. The neuronal morphology was comparable to neurons co-cultured with rodent astrocytes^12,70–73^, showing fine dendritic outgrowth and relatively small nuclei, with no neuronal clustering. Although we observed healthy astrocytes for all different co-cultures, some morphological differences of the astrocytes were observed between hiPSC-lines and batches. For example, the astrocytes of hiPSC-line 1 in batch 2 showed a heterogenous mix of naïve and potentially reactive (star-shaped) astrocytes, which we therefore consider a co-culture of excellent quality. In contrast to the astrocytes of hiPSC-line 1 in batch 1, which are relatively flat and morphologically very similar to most (DIV 42) monoculture astrocytes, the astrocytes in the second batch clearly show increased star-shaped morphology (as indicated by an increased number of astrocytic protrusions), compared to monocultures. However, the differences in astrocytic morphology did not seem to affect neuronal morphology between the two batches. Interestingly, a lower survival rate of astrocytes also did not necessarily result in decreased neuronal survival or quality, as shown by the co-cultures with astrocytes of hiPSC-line 7. hiPSC-line 7 showed lower astrocyte survival in both batches indicating that this could be line specific, and possibly solved by plating a higher number of astrocytes to compensate.

**Figure 5.**
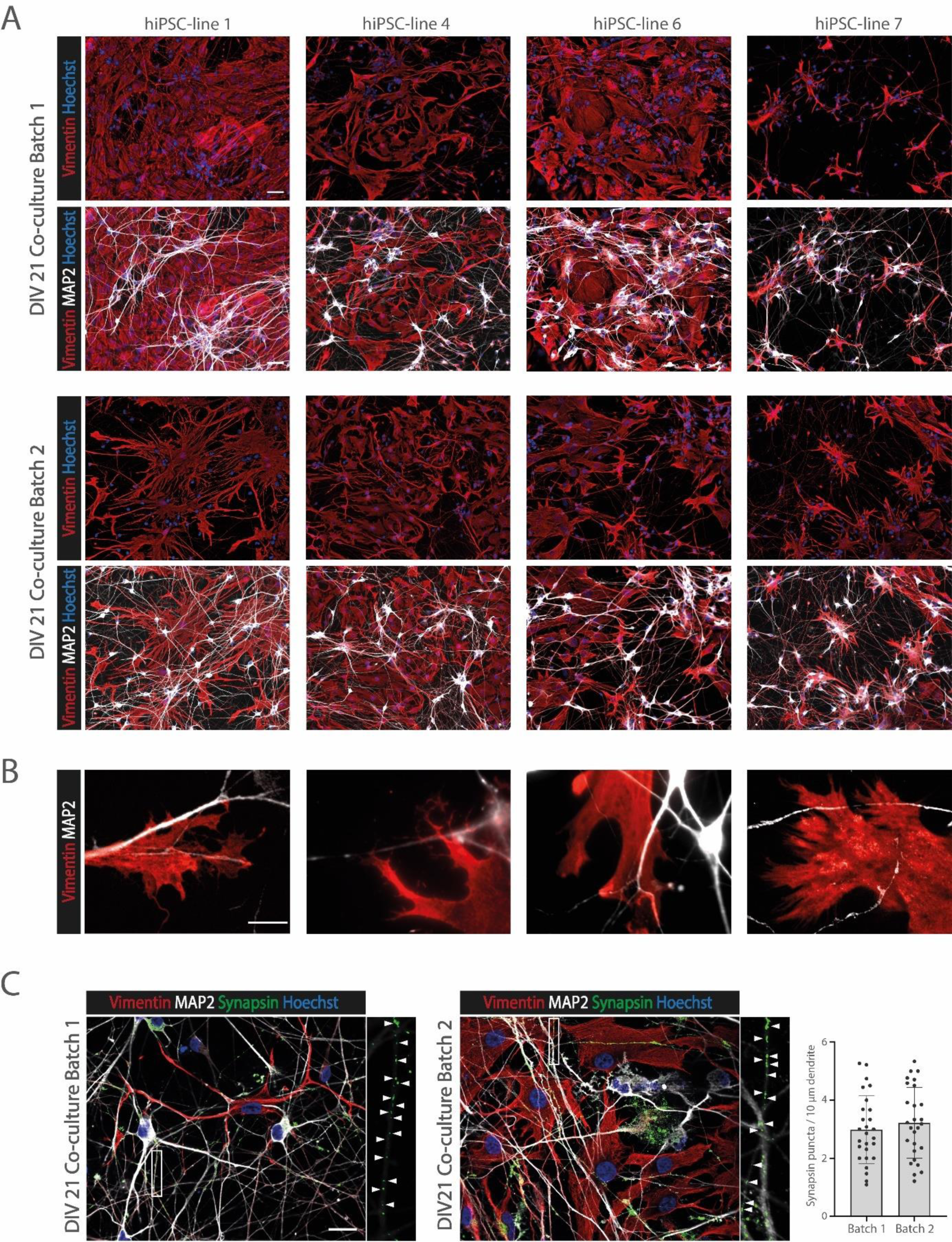
Immunofluorescence of hiPSC-astrocyte and iNeuron co-cultures. **(A)** Immunostaining of Vimentin (red) and MAP2 (white) of two independent batches of DIV 21 co-cultures with iNeurons (hiPSC-line 1) together with hiPSC-astrocytes generated from hiPSC-line 1, hiPSC-line 4, hiPSC-line 6 and hiPSC-line 7. All pictures were taken at the same magnification (scale bar = 50 µm). **(B)** Immunostaining of Vimentin (red) and MAP2 (white) of DIV 21 co-cultures with iNeurons (hiPSC-line 1) together with hiPSC-astrocytes generated from hiPSC-line 1, showing interaction of the astrocytic end feet with the neuronal dendrites. All pictures were taken at the same magnification (scale bar = 10 µm). **(C)** Immunostaining of Vimentin (red), MAP2 (white) and synapsin (green) of two independent batches of DIV 21 co-cultures with iNeurons (hiPSC-line 1) together with hiPSC-astrocytes generated from hiPSC-line 4. All pictures were taken at the same magnification (scale bar = 20 µm). White arrows indicate synapsin puncta. One-way ANOVA of the number of synapsin puncta per 10 µm neuronal dendrite was not significantly different between the two independent co-culture batches (n= 15 neurons).

In addition, we observed arborizations of the astrocytic end feet, interacting with their surrounding neurons (Figure 5B). To investigate whether the hiPSC-astrocytes also support synapse formation, we quantified the number of synapses within the co-cultures using immunofluorescent labelling of Synapsin (Figure 5C). We observed no significant differences in the number of Synapsin puncta between two different co-culture batches. The number of Synapsin puncta (∼3 puncta/10 µm), are in the same range as observed for the standard co-cultures with rodent astrocytes^70–73^.

### hiPSC-astrocytes support robust synchronization of the neuronal activity of iNeurons on MEA

Immunofluorescent imaging was used to confirm that the hiPSC-astrocytes support neuronal development, maturation as well as synapse formation when co-cultured with iNeurons. As described previously, to assess whether the iNeurons in these co-cultures also form synchronized neuronal networks, MEAs were used to measure spontaneous neuronal activity. Extensive testing of numerous adjustments to the protocol (Supplementary Figure 7), assessed using the MEAs, ultimately led to the development of a successful co-culture approach (see Methods section for details). In summary, Ngn2 stable hiPSCs were differentiated towards neurons by addition of doxycycline. In contrast to the protocol using rodent astrocytes, hiPSC-astrocytes were added at DIV 6 (of the neuronal differentiation), in a 2:1 ratio instead of 1:1. Additionally, at DIV 6 the cells were cultured in the presence of RevitaCell^TM^ to promote hiPSC-astrocyte survival and with AM FCS instead of the standard FCS used previously (Figure 6A).

**Figure 6.**
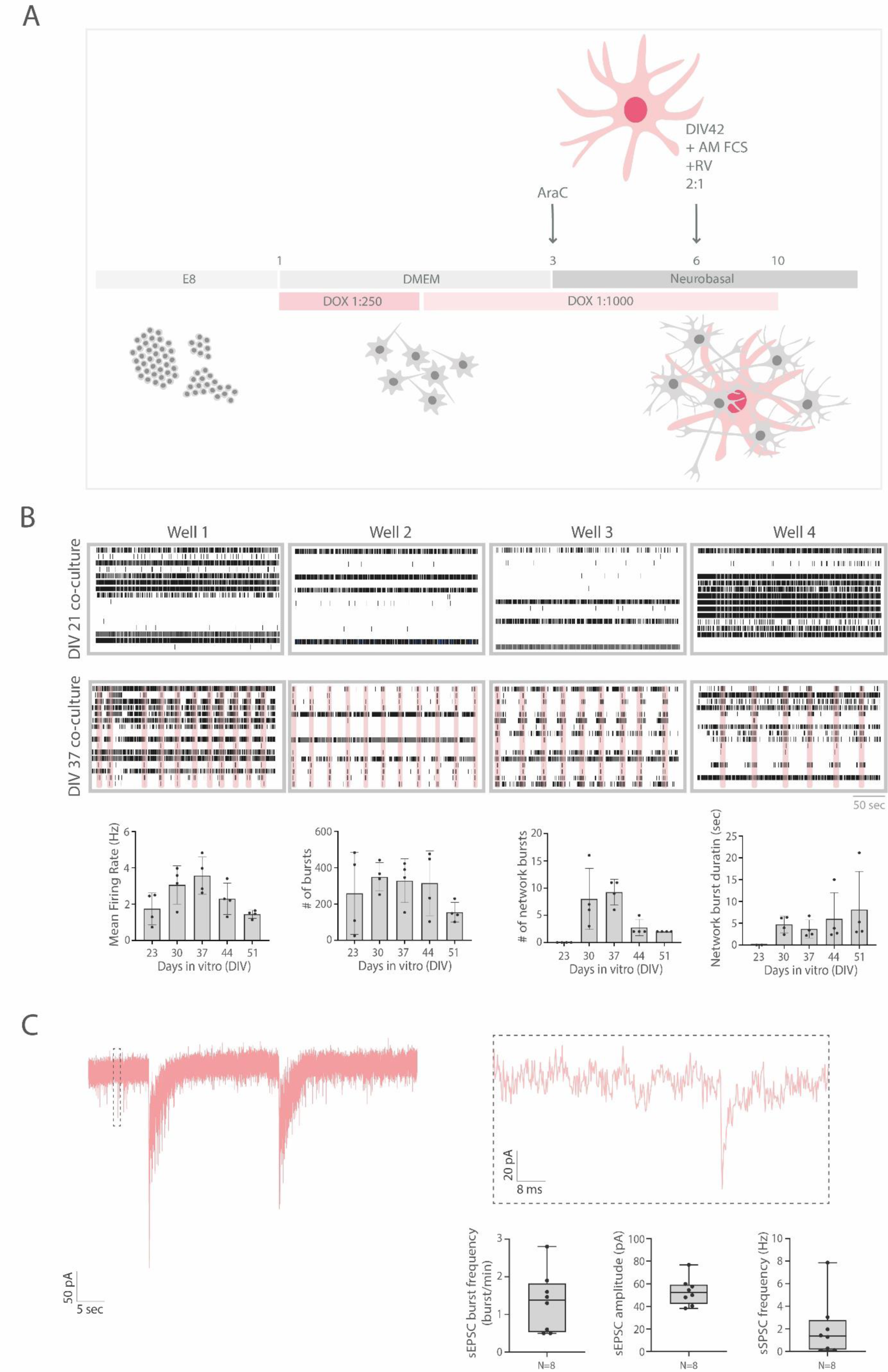
MEA analysis confirming neuronal network activity in hiPSC-astrocyte and iNeuron co-cultures. **(A)** Schematic representation of the protocol to co-culture hiPSC-astrocytes with iNeurons. **(B)** Raster plots of the four different wells of the DIV 23 and DIV 37 co-culture with iNeurons (hiPSC-line 1) together with hiPSC-astrocytes (scale bar = 50 seconds). All four wells show consistent network burst frequency at DIV 37. In addition, mean firing rate, number of bursts, number of network bursts and network burst duration of the four wells at DIV 21, 30, 37, 44 and 51 is shown. **(D)** Representative 60-sec whole-cell voltage-clamp recording of spontaneous excitatory postsynaptic currents (sEPSCs) of DIV 37 co-culture with iNeurons (hiPSC-line 1) together with hiPSC-astrocytes, with 90 ms zoom-in of an sEPSC. Quantification of sEPSC burst frequency, sEPSC amplitude and sEPSC frequency (n = 8) is shown.

Using the final co-culture protocol, the iNeurons in all four biological replicates showed a consistent number of NBs and NB duration (Figure 6B). Although the NB frequency changed over time, similarly as described for co-cultures with rodent astrocytes, we observed that the number of NBs within the biological replicates for each time point is highly consistent. Other parameters such as the number of bursts and the mean firing rate show higher variability within biological replicates. However, some of these parameters are generally considered to be more variable across biological replicates^73^. Interestingly, the network activity of the co-cultures with hiPSC-astrocytes is delayed (starting at DIV 30) compared to co-cultures with rodent astrocytes (starting at DIV 16/23). This is in line with studies showing delayed neuronal development in humans compared to rodents and the great apes^74–78^. Finally, using whole-cell voltage-clamp recordings we recorded spontaneous excitatory synaptic currents at DIV35. All recorded neurons received synaptic input and showed burst activity (Figure 6C).

## Discussion

We developed a protocol for hiPSC-derived astrocytes which can support neuronal development when co-cultured together with iNeurons. In contrast to other protocols which either use ectopic gene expression or intermediate NPC stages, our protocol directly differentiates hiPSCs towards functional astrocytes using only commercial AM. The protocol has been validated in thirty different hiPSC-lines to varying extents. Here we have described a detailed characterization of a subset of these hiPSC-astrocytes cultures both in monoculture as well as in co-culture together with iNeurons.

### hiPSC-astrocytes in monoculture

Morphological differences defined by astrocyte volume, morphological complexity and proliferation rate were observed between the different hiPSC-lines, both across different differentiation batches (Supplementary Figure 2 and 3) as well as within batches (Figure 1C). Nevertheless, these morphological differences are consistent for each hiPSC-line across different batches, demonstrating minimal batch-to-batch variability (Supplementary Figure 4). So the morphological differences between hiPSC-lines most likely originate from genetic heterogeneity rather than environmental factors. As mentioned before, a potential factor influencing differentiation efficiency of the astrocytes (as immature astrocytes tend to grow faster than mature astrocytes) could be the method used for reprogramming of the hiPSC-lines. Differences in reprogramming efficiency between the different vectors (Sendai, episomal and retroviral vectors) are known^79–81^. For example, reprogramming efficiency was shown to be less efficient using retroviral vectors compared to episomal and Sendai virus vectors^82^, which could potentially also affect astrocyte differentiation efficiency. However, additional research is needed to further investigate this.

Due to the significance of astrocytic diversity in the brain^83^, it was anticipated that heterogenous hiPSC-astrocyte cultures would yield superior astrocytes compared to more homogeneous hiPSC-astrocyte cultures. This superiority could manifest in various ways, such as the expression of a wider array of astrocytic genes or the secretion of more astrocyte-specific proteins. Surprisingly, no discernible correlation between the heterogeneity of the astrocyte monocultures (or other morphological differences) and the functionality or quality of the co-cultures was observed. Despite the acknowledged importance of astrocytic diversity in *in vivo* brain development^83^, the limited effect of the astrocyte monoculture heterogeneity on the co-culture quality suggests that such diversity may not be as crucial for *in vitro* neuronal network development. However, in this study the differences were mainly based on morphological heterogeneity and proliferation rate. Conducting more comprehensive analyses, such as single-cell RNA sequencing, could potentially elucidate the correlation between these morphological disparities and astrocytic function.

### hiPSC-astrocytes in co-culture with iNeurons

We also assessed the ability of the hiPSC-astrocytes to support neuronal (network) development when co-cultured with iNeurons. Considering that these protocols are developed for disease modelling and phenotyping, our most important criteria were robustness and reproducibility. As starting point our “standard” co-culture protocol^12^ was tested, using immunofluorescent imaging and MEA as the main readouts to assess the quality and functionality of the co-cultures. Upon testing many adjusted versions of the protocol, the co-culture protocol described in the methods section is currently the most optimal for both the hiPSC-astrocytes and iNeurons and considered reproducible. Some of these adjustments significantly improved the quality of the co-cultures. One of them was the use of FBS from the AM kit instead of regular FBS, most likely due to the lower concentrations of endotoxins. Moreover, addition of FBS to the medium at the moment of astrocyte plating instead of 7 days after astrocyte plating, significantly improved the viability of the hiPSC-astrocytes in co-culture. DIV 42 was shown to be the most optimal timepoint for the hiPSC-astrocytes to co-mature with the iNeurons. However, given the differences between the hiPSC-lines during astrocyte differentiation, the most optimal timepoint of plating might vary between the different hiPSC-lines. Lastly, due to the increased cell size of the hiPSC-astrocytes compared to rodent astrocytes in monoculture, possibly resulting in increased stress due to limited space, different ratios of neurons to astrocytes were assessed. Changing the 1:1 neuron to astrocyte ratio into 2:1 ratio greatly improved the quality of the co-cultures.

In contrast to other *in vitro* studies^15,20^, the number of synapses in co-cultures with hiPSC-astrocytes is similar to co-cultures with rodent astrocytes. We also demonstrated that when co-cultured with hiPSC-astrocytes, neuronal network activity started around DIV 30, while previous research has shown that when co-cultured with *ex vivo* rodent astrocytes, neuronal network activity starts around DIV 14^73^. In line with literature, this may suggest that human brain development compared to rodent brain development is delayed. However, in order to investigate the effect of astrocytes on the number of synapses or neuronal network activity in rodents versus humans (which is beyond the scope of this study), *ex vivo* primary astrocytes should not be compared to *in vitro* hiPSC-derived astrocytes. Especially since neurons and other glial cell types highly influence astrocyte development, both via direct contact^84,85^ but also indirectly by the secretion of metabolites and proteins^84–88^. Therefore, to assess potential differences between human and rodent astrocytes, either human and rodent iPSC-astrocytes, or *ex vivo* primary human and rodent astrocytes should be used.

Although the morphological differences between different batches of the astrocyte monocultures were minimal, significant differences in astrocytic morphology were observed between different batches of co-cultures using the same hiPSC-line. In one batch the hiPSC-astrocytes co-matured into a heterogenous mix of naïve and potentially reactive (star-shaped) astrocytes, while in the other batch the hiPSC-astrocytes appeared to fail to co-mature and therefore maintained their flat monoculture-like morphology. It is still unclear how this difference can be explained, but it most likely originates from variables during the hiPSC-astrocyte differentiation, including the procedure of splitting the astrocytes during differentiation. Although the procedure might seem consistent, the difference between splitting a 95% confluent flask at 1:3 ratio compared to splitting a 100% confluent flask in 1:3 ratio, might have had more impact than previously anticipated. Especially given that confluency, cell density and cell-cell contact greatly impact glial cell development^89^. In order to improve consistency; instead of splitting at 1:3 ratio, a specific number of cells e.g., 1.000.000 per T75 flask could be re-seeded. Therefore, the cell number might be more consistent during differentiation, and across different hiPSC-astrocyte batches. Surprisingly, the flat monoculture-like astrocytes seemed to support the neurons as well as the heterogeneous astrocyte cultures including the star-shaped astrocytes. This indicates that in co-culture, the differences in astrocyte morphology defined by e.g., the number of astrocytic protrusions, does not affect neuronal growth and maturation which may suggest that so-called naïve astrocytes are of more importance in *in vitro* co-cultures compared to the reactive star-shaped astrocytes^32^. However, here the quality of the neurons was solely assessed based on morphology, so potential differences e.g., at gene expression or electrophysiological level should be identified in additional studies.

### The effect of skipping the NPC intermediate stage during astrocyte differentiation

As mentioned before, most *in vitro* hiPSC-astrocyte differentiation protocols first differentiate the hiPSCs towards NPCs, which later are differentiated towards astrocytes. Since this protocol does not include an intermediate NPC differentiation step, we compared the hiPSC-astrocytes to astrocytes derived from NPCs (NPC-astrocytes) using bulk RNA sequencing. Interestingly, hiPSC-astrocytes and NPC-astrocytes clustered closely together, confirming minimal differences between these two protocols. Recently, Räsänen et al. (2024) also demonstrated neuronal network activity in hiPSC-astrocyte and iNeuron co-cultures, showing a similar developmental trajectory of the human co-culture over time as was shown in this study^90^. This further indicates that skipping of the NPC intermediate stage has minimal effect on *in vitro* astrocyte development, and subsequent neuronal network development. Although we did not specifically differentiate the hiPSCs towards NPCs using e.g., dual SMAD inhibition, it could still be that using the commercial AM (of which the exact composition is unknown) the hiPSCs first differentiate towards a NPC-like stage or astrocyte precursor cell type and later towards functional astrocytes. Since the astrocyte differentiation protocol used in this study is based on a commercial AM kit, the effect of potential changes of the medium by the manufacturer are unknown. The risk of batch-to-batch variability due to potential differences in the medium (or FBS) are therefore also increased.

In conclusion, our results show that direct differentiation of hiPSCs into astrocytes using commercial AM results consistently in mature and functional astrocytes. These hiPSC-astrocytes can functionally support neuronal network formation when co-cultured with iNeurons showing consistent and robust neuronal network patterns. Although variability between different hiPSC-lines is acknowledged, batch-to-batch variability is minimal. The results from thirty hiPSC lines have shown that both the hiPSC-astrocyte differentiation protocol as well as the co-culture protocol are robust and consistent allowing it to be used for cell-type specific disease modelling and phenotyping.

As mentioned before, we see this work as a living document, and we would like to encourage those who are interested, to test these protocols within their own labs using their own hiPSC lines and subsequently share their data with us (please contact the corresponding authors) to further validate and improve these protocols which we will jointly publish in the 2.0 version of this manuscript.

## Acknowledgments

We gratefully acknowledge Dr. Shan Wang, Dr. Brooke Latour, Carlos González Jiménez and Mara Graziani for the stimulating discussions. We also want to acknowledge the members of the Changing rare disorders of lysine metabolism (CHARLIE) consortium (EJPRD grant nr: 825575 awarded to C.v.K. and N.N.K.). I.M.E.S is supported by an internal Radboudumc PhD grant provided by the Radboud Institute for Molecular Life Sciences to C.v.K and A.G.

## References

1. Somjen, G.G. (1988). Nervenkitt: notes on the history of the concept of neuroglia. Glia 1, 2–9. 10.1002/glia.440010103.

2. He, F., and Sun, Y.E. (2007). Glial cells more than support cells? Int J Biochem Cell Biol 39, 661–665. 10.1016/j.biocel.2006.10.022.

3. Volterra, A., and Meldolesi, J. (2005). Astrocytes, from brain glue to communication elements: the revolution continues. Nat Rev Neurosci 6, 626–640. 10.1038/nrn1722.

4. Sidoryk-Wegrzynowicz, M., Wegrzynowicz, M., Lee, E., Bowman, A.B., and Aschner, M. (2011). Role of astrocytes in brain function and disease. Toxicol Pathol 39, 115–123. 10.1177/0192623310385254.

5. Siracusa, R., Fusco, R., and Cuzzocrea, S. (2019). Astrocytes: Role and Functions in Brain Pathologies. Front Pharmacol 10, 1114. 10.3389/fphar.2019.01114.

6. Kim, Y., Park, J., and Choi, Y.K. (2019). The Role of Astrocytes in the Central Nervous System Focused on BK Channel and Heme Oxygenase Metabolites: A Review. Antioxidants (Basel) 8. 10.3390/antiox8050121.

7. Pietilainen, O., Trehan, A., Meyer, D., Mitchell, J., Tegtmeyer, M., Valakh, V., Gebre, H., Chen, T., Vartiainen, E., Farhi, S.L., et al. (2023). Astrocytic cell adhesion genes linked to schizophrenia correlate with synaptic programs in neurons. Cell Rep 42, 111988. 10.1016/j.celrep.2022.111988.

8. Koskuvi, M., Lehtonen, S., Trontti, K., Keuters, M., Wu, Y.C., Koivisto, H., Ludwig, A., Plotnikova, L., Virtanen, P.L.J., Rasanen, N., et al. (2022). Contribution of astrocytes to familial risk and clinical manifestation of schizophrenia. Glia 70, 650–660. 10.1002/glia.24131.

9. Jantti, H., Oksanen, M., Kettunen, P., Manta, S., Mouledous, L., Koivisto, H., Ruuth, J., Trontti, K., Dhungana, H., Keuters, M., et al. (2022). Human PSEN1 Mutant Glia Improve Spatial Learning and Memory in Aged Mice. Cells 11. 10.3390/cells11244116.

10. Di Giorgio, F.P., Carrasco, M.A., Siao, M.C., Maniatis, T., and Eggan, K. (2007). Non-cell autonomous effect of glia on motor neurons in an embryonic stem cell-based ALS model. Nat Neurosci 10, 608–614. 10.1038/nn1885.

11. Astick, M., and Vanderhaeghen, P. (2018). From Human Pluripotent Stem Cells to Cortical Circuits. Curr Top Dev Biol 129, 67–98. 10.1016/bs.ctdb.2018.02.011.

12. Frega, M., van Gestel, S.H., Linda, K., van der Raadt, J., Keller, J., Van Rhijn, J.R., Schubert, D., Albers, C.A., and Nadif Kasri, N. (2017). Rapid Neuronal Differentiation of Induced Pluripotent Stem Cells for Measuring Network Activity on Micro-electrode Arrays. J Vis Exp. 10.3791/54900.

13. Lin, H.C., He, Z., Ebert, S., Schornig, M., Santel, M., Nikolova, M.T., Weigert, A., Hevers, W., Kasri, N.N., Taverna, E., et al. (2021). NGN2 induces diverse neuron types from human pluripotency. Stem Cell Reports 16, 2118–2127. 10.1016/j.stemcr.2021.07.006.

14. Zhang, Y., Pak, C., Han, Y., Ahlenius, H., Zhang, Z., Chanda, S., Marro, S., Patzke, C., Acuna, C., Covy, J., et al. (2013). Rapid single-step induction of functional neurons from human pluripotent stem cells. Neuron 78, 785–798. 10.1016/j.neuron.2013.05.029.

15. Vasile, F., Dossi, E., and Rouach, N. (2017). Human astrocytes: structure and functions in the healthy brain. Brain Struct Funct 222, 2017–2029. 10.1007/s00429-017-1383-5.

16. Perriot, S., Canales, M., Mathias, A., and Du Pasquier, R. (2021). Differentiation of functional astrocytes from human-induced pluripotent stem cells in chemically defined media. STAR Protoc 2, 100902. 10.1016/j.xpro.2021.100902.

17. Tcw, J., Wang, M., Pimenova, A.A., Bowles, K.R., Hartley, B.J., Lacin, E., Machlovi, S.I., Abdelaal, R., Karch, C.M., Phatnani, H., et al. (2017). An Efficient Platform for Astrocyte Differentiation from Human Induced Pluripotent Stem Cells. Stem Cell Reports 9, 600–614. 10.1016/j.stemcr.2017.06.018.

18. Mulica, P., Venegas, C., Landoulsi, Z., Badanjak, K., Delcambre, S., Tziortziou, M., Hezzaz, S., Ghelfi, J., Smajic, S., Schwamborn, J., et al. (2023). Comparison of two protocols for the generation of iPSC-derived human astrocytes. Biol Proced Online 25, 26. 10.1186/s12575-023-00218-x.

19. Chandrasekaran, A., Avci, H.X., Leist, M., Kobolak, J., and Dinnyes, A. (2016). Astrocyte Differentiation of Human Pluripotent Stem Cells: New Tools for Neurological Disorder Research. Front Cell Neurosci 10, 215. 10.3389/fncel.2016.00215.

20. B. Lendemeijer, M.U., B. Mossink, S. Hijazi, S.G. Sampedro, G. Shpak, D.E. Slump M.C.G.N., van den Hout, W.F.J.v.I., E.M.J. Bindels, W.J.G. Hoogendijk, N. Nadif Kasri, F.M.S. de Vrij, and Kushner, S.A. (2022). Rapid specification of human pluripotent stem cells to functional astrocytes BioRxiv. 10.1101/2022.08.25.505166.

21. Voulgaris, D., Nikolakopoulou, P., and Herland, A. (2022). Generation of Human iPSC-Derived Astrocytes with a mature star-shaped phenotype for CNS modeling. Stem Cell Rev Rep 18, 2494–2512. 10.1007/s12015-022-10376-2.

22. de Leeuw, V.C., van Oostrom, C.T.M., Westerink, R.H.S., Piersma, A.H., Heusinkveld, H.J., and Hessel, E.V.S. (2020). An efficient neuron-astrocyte differentiation protocol from human embryonic stem cell-derived neural progenitors to assess chemical-induced developmental neurotoxicity. Reprod Toxicol 98, 107–116. 10.1016/j.reprotox.2020.09.003.

23. Shaltouki, A., Peng, J., Liu, Q., Rao, M.S., and Zeng, X. (2013). Efficient generation of astrocytes from human pluripotent stem cells in defined conditions. Stem Cells 31, 941–952. 10.1002/stem.1334.

24. Jovanovic, V.M., Weber, C., Slamecka, J., Ryu, S., Chu, P.H., Sen, C., Inman, J., De Sousa, J.F., Barnaeva, E., Hirst, M., et al. (2023). A defined roadmap of radial glia and astrocyte differentiation from human pluripotent stem cells. Stem Cell Reports 18, 1701–1720. 10.1016/j.stemcr.2023.06.007.

25. Li, X., Tao, Y., Bradley, R., Du, Z., Tao, Y., Kong, L., Dong, Y., Jones, J., Yan, Y., Harder, C.R.K., et al. (2018). Fast Generation of Functional Subtype Astrocytes from Human Pluripotent Stem Cells. Stem Cell Reports 11, 998–1008. 10.1016/j.stemcr.2018.08.019.

26. Neyrinck, K., Van Den Daele, J., Vervliet, T., De Smedt, J., Wierda, K., Nijs, M., Vanbokhoven, T., D’Hondt, A., Planque, M., Fendt, S.M., et al. (2021). SOX9-induced Generation of Functional Astrocytes Supporting Neuronal Maturation in an All-human System. Stem Cell Rev Rep 17, 1855–1873. 10.1007/s12015-021-10179-x.

27. Soubannier, V., Maussion, G., Chaineau, M., Sigutova, V., Rouleau, G., Durcan, T.M., and Stifani, S. (2020). Characterization of human iPSC-derived astrocytes with potential for disease modeling and drug discovery. Neurosci Lett 731, 135028. 10.1016/j.neulet.2020.135028.

28. Yeon, G.B., Shin, W.H., Yoo, S.H., Kim, D., Jeon, B.M., Park, W.U., Bae, Y., Park, J.Y., You, S., Na, D., and Kim, D.S. (2021). NFIB induces functional astrocytes from human pluripotent stem cell-derived neural precursor cells mimicking in vivo astrogliogenesis. J Cell Physiol 236, 7625–7641. 10.1002/jcp.30405.

29. Westergard, T., and Rothstein, J.D. (2020). Astrocyte Diversity: Current Insights and Future Directions. Neurochem Res 45, 1298–1305. 10.1007/s11064-020-02959-7.

30. Ben Haim, L., and Rowitch, D.H. (2017). Functional diversity of astrocytes in neural circuit regulation. Nat Rev Neurosci 18, 31–41. 10.1038/nrn.2016.159.

31. Khakh, B.S., and Sofroniew, M.V. (2015). Diversity of astrocyte functions and phenotypes in neural circuits. Nat Neurosci 18, 942–952. 10.1038/nn.4043.

32. Gottipati, M.K., Zuidema, J.M., and Gilbert, R.J. (2020). Biomaterial strategies for creating in vitro astrocyte cultures resembling in vivo astrocyte morphologies and phenotypes. Curr Opin Biomed Eng 14, 67–74. 10.1016/j.cobme.2020.06.004.

33. Stifani, S. (2021). Taking Cellular Heterogeneity Into Consideration When Modeling Astrocyte Involvement in Amyotrophic Lateral Sclerosis Using Human Induced Pluripotent Stem Cells. Front Cell Neurosci 15, 707861. 10.3389/fncel.2021.707861.

34. Schuurmans, I.M.E., Wu, K.M., van Karnebeek, C.D.M., Nadif Kasri, N., and Garanto, A. (2023). Generation of an induced pluripotent stem cell line carrying biallelic deletions (SCTCi019-B) in ALDH7A1 using CRISPR/Cas9. Stem Cell Res 71, 103173. 10.1016/j.scr.2023.103173.

35. Schuurmans, I.M.E., Wu, K.M., van Karnebeek, C.D.M., Nadif Kasri, N., and Garanto, A. (2023). Generation of an induced pluripotent stem cell line carrying a biallelic deletion (SCTCi019-A) in GCDH using CRISPR/Cas9. Stem Cell Res 69, 103069. 10.1016/j.scr.2023.103069.

36. Dyke, E., Bijnagte-Schoenmaker, C., Wu, K.M., Oudakker, A., Roepman, R., and Nadif Kasri, N. (2023). Generation of induced pluripotent stem cell line carrying frameshift variants in NPHP1 (UCSFi001-A-68) using CRISPR/Cas9. Stem Cell Res 68, 103053. 10.1016/j.scr.2023.103053.

37. Shi, Y., Kirwan, P., and Livesey, F.J. (2012). Directed differentiation of human pluripotent stem cells to cerebral cortex neurons and neural networks. Nat Protoc 7, 1836–1846. 10.1038/nprot.2012.116.

38. Frega, M., Linda, K., Keller, J.M., Gumus-Akay, G., Mossink, B., van Rhijn, J.R., Negwer, M., Klein Gunnewiek, T., Foreman, K., Kompier, N., et al. (2019). Neuronal network dysfunction in a model for Kleefstra syndrome mediated by enhanced NMDAR signaling. Nat Commun 10, 4928. 10.1038/s41467-019-12947-3.

39. Chen, S., Zhou, Y., Chen, Y., and Gu, J. (2018). fastp: an ultra-fast all-in-one FASTQ preprocessor. Bioinformatics 34, i884–i890. 10.1093/bioinformatics/bty560.

40. Li, H., Handsaker, B., Wysoker, A., Fennell, T., Ruan, J., Homer, N., Marth, G., Abecasis, G., Durbin, R., and Genome Project Data Processing, S. (2009). The Sequence Alignment/Map format and SAMtools. Bioinformatics 25, 2078-2079. 10.1093/bioinformatics/btp352.

41. Smith, T., Heger, A., and Sudbery, I. (2017). UMI-tools: modeling sequencing errors in Unique Molecular Identifiers to improve quantification accuracy. Genome Res 27, 491–499. 10.1101/gr.209601.116.

42. Liao, Y., Smyth, G.K., and Shi, W. (2014). featureCounts: an efficient general purpose program for assigning sequence reads to genomic features. Bioinformatics 30, 923–930. 10.1093/bioinformatics/btt656.

43. Zhang, Y., Sloan, S.A., Clarke, L.E., Caneda, C., Plaza, C.A., Blumenthal, P.D., Vogel, H., Steinberg, G.K., Edwards, M.S., Li, G., et al. (2016). Purification and Characterization of Progenitor and Mature Human Astrocytes Reveals Transcriptional and Functional Differences with Mouse. Neuron 89, 37–53. 10.1016/j.neuron.2015.11.013.

44. Ritchie, M.E., Phipson, B., Wu, D., Hu, Y., Law, C.W., Shi, W., and Smyth, G.K. (2015). limma powers differential expression analyses for RNA-sequencing and microarray studies. Nucleic Acids Res 43, e47. 10.1093/nar/gkv007.

45. van Hugte, E.J.H., Lewerissa, E.I., Wu, K.M., Scheefhals, N., Parodi, G., van Voorst, T.W., Puvogel, S., Kogo, N., Keller, J.M., Frega, M., et al. (2023). SCN1A-deficient excitatory neuronal networks display mutation-specific phenotypes. Brain 146, 5153–5167. 10.1093/brain/awad245.

46. Marino, M., Misuri, L., and Brogioli, D. (2014). A new open source software for the calculation of the liquid junction potential between two solutions according to the stationary Nernst-Planck equation. arXiv preprint arXiv:1403.3640.

47. Draberova, E., Del Valle, L., Gordon, J., Markova, V., Smejkalova, B., Bertrand, L., de Chadarevian, J.P., Agamanolis, D.P., Legido, A., Khalili, K., et al. (2008). Class III beta-tubulin is constitutively coexpressed with glial fibrillary acidic protein and nestin in midgestational human fetal astrocytes: implications for phenotypic identity. J Neuropathol Exp Neurol 67, 341–354. 10.1097/NEN.0b013e31816a686d.

48. Khakh, B.S., and Deneen, B. (2019). The Emerging Nature of Astrocyte Diversity. Annu Rev Neurosci 42, 187–207. 10.1146/annurev-neuro-070918-050443.

49. Endo, F., Kasai, A., Soto, J.S., Yu, X., Qu, Z., Hashimoto, H., Gradinaru, V., Kawaguchi, R., and Khakh, B.S. (2022). Molecular basis of astrocyte diversity and morphology across the CNS in health and disease. Science 378, eadc9020. 10.1126/science.adc9020.

50. Barbar, L., Jain, T., Zimmer, M., Kruglikov, I., Sadick, J.S., Wang, M., Kalpana, K., Rose, I.V.L., Burstein, S.R., Rusielewicz, T., et al. (2020). CD49f Is a Novel Marker of Functional and Reactive Human iPSC-Derived Astrocytes. Neuron 107, 436–453 e412. 10.1016/j.neuron.2020.05.014.

51. Nielsen, S., Nagelhus, E.A., Amiry-Moghaddam, M., Bourque, C., Agre, P., and Ottersen, O.P. (1997). Specialized membrane domains for water transport in glial cells: high-resolution immunogold cytochemistry of aquaporin-4 in rat brain. J Neurosci 17, 171–180. 10.1523/JNEUROSCI.17-01-00171.1997.

52. Frigeri, A., Gropper, M.A., Umenishi, F., Kawashima, M., Brown, D., and Verkman, A.S. (1995). Localization of MIWC and GLIP water channel homologs in neuromuscular, epithelial and glandular tissues. J Cell Sci 108 (Pt 9), 2993–3002. 10.1242/jcs.108.9.2993.

53. Mader, S., and Brimberg, L. (2019). Aquaporin-4 Water Channel in the Brain and Its Implication for Health and Disease. Cells 8. 10.3390/cells8020090.

54. Boal, A.M., Risner, M.L., Cooper, M.L., Wareham, L.K., and Calkins, D.J. (2021). Astrocyte Networks as Therapeutic Targets in Glaucomatous Neurodegeneration. Cells 10. 10.3390/cells10061368.

55. Cooper, M.L., Pasini, S., Lambert, W.S., D’Alessandro, K.B., Yao, V., Risner, M.L., and Calkins, D.J. (2020). Redistribution of metabolic resources through astrocyte networks mitigates neurodegenerative stress. Proc Natl Acad Sci U S A 117, 18810–18821. 10.1073/pnas.2009425117.

56. Wang, X., Takano, T., and Nedergaard, M. (2009). Astrocytic calcium signaling: mechanism and implications for functional brain imaging. Methods Mol Biol 489, 93–109. 10.1007/978-1-59745-543-5_5.

57. Chung, W.S., Allen, N.J., and Eroglu, C. (2015). Astrocytes Control Synapse Formation, Function, and Elimination. Cold Spring Harb Perspect Biol 7, a020370. 10.1101/cshperspect.a020370.

58. Jha, M.K., Kim, J.H., Song, G.J., Lee, W.H., Lee, I.K., Lee, H.W., An, S.S.A., Kim, S., and Suk, K. (2018). Functional dissection of astrocyte-secreted proteins: Implications in brain health and diseases. Prog Neurobiol 162, 37–69. 10.1016/j.pneurobio.2017.12.003.

59. Okabe, K., Fukada, H., Tai-Nagara, I., Ando, T., Honda, T., Nakajima, K., Takeda, N., Fong, G.H., Ema, M., and Kubota, Y. (2020). Neuron-derived VEGF contributes to cortical and hippocampal development independently of VEGFR1/2-mediated neurotrophism. Dev Biol 459, 65–71. 10.1016/j.ydbio.2019.11.016.

60. Yepes, M. (2007). TWEAK and the central nervous system. Mol Neurobiol 35, 255–265. 10.1007/s12035-007-0024-z.

61. Desole, C., Gallo, S., Vitacolonna, A., Montarolo, F., Bertolotto, A., Vivien, D., Comoglio, P., and Crepaldi, T. (2021). HGF and MET: From Brain Development to Neurological Disorders. Front Cell Dev Biol 9, 683609. 10.3389/fcell.2021.683609.

62. Erta, M., Quintana, A., and Hidalgo, J. (2012). Interleukin-6, a major cytokine in the central nervous system. Int J Biol Sci 8, 1254–1266. 10.7150/ijbs.4679.

63. Yao, Y., and Tsirka, S.E. (2014). Monocyte chemoattractant protein-1 and the blood-brain barrier. Cell Mol Life Sci 71, 683–697. 10.1007/s00018-013-1459-1.

64. Filipovic, R., and Zecevic, N. (2008). The effect of CXCL1 on human fetal oligodendrocyte progenitor cells. Glia 56, 1–15. 10.1002/glia.20582.

65. Easley-Neal, C., Foreman, O., Sharma, N., Zarrin, A.A., and Weimer, R.M. (2019). CSF1R Ligands IL-34 and CSF1 Are Differentially Required for Microglia Development and Maintenance in White and Gray Matter Brain Regions. Front Immunol 10, 2199. 10.3389/fimmu.2019.02199.

66. Rose-John, S. (2018). Interleukin-6 Family Cytokines. Cold Spring Harb Perspect Biol 10. 10.1101/cshperspect.a028415.

67. Bonni, A., Sun, Y., Nadal-Vicens, M., Bhatt, A., Frank, D.A., Rozovsky, I., Stahl, N., Yancopoulos, G.D., and Greenberg, M.E. (1997). Regulation of gliogenesis in the central nervous system by the JAK-STAT signaling pathway. Science 278, 477–483. 10.1126/science.278.5337.477.

68. Fukuda, S., Abematsu, M., Mori, H., Yanagisawa, M., Kagawa, T., Nakashima, K., Yoshimura, A., and Taga, T. (2007). Potentiation of astrogliogenesis by STAT3-mediated activation of bone morphogenetic protein-Smad signaling in neural stem cells. Mol Cell Biol 27, 4931–4937. 10.1128/MCB.02435-06.

69. Marta Cerina, M.C.P.a.M.F. (2023). The potential of in vitro neuronal networks cultured on micro electrode arrays for biomedical research. Progress in Biomedical Engineering 5. 10.1088/2516-1091/acce12.

70. Mijdam, R., Bijnagte-Schoenmaker, C., Dyke, E., Moons, S.J., Boltje, T.J., Nadif Kasri, N., and Lefeber, D.J. (2023). Sialic acid biosynthesis pathway blockade disturbs neuronal network formation in human iPSC-derived excitatory neurons. J Neurochem 167, 76–89. 10.1111/jnc.15934.

71. Klein Gunnewiek, T.M., Van Hugte, E.J.H., Frega, M., Guardia, G.S., Foreman, K., Panneman, D., Mossink, B., Linda, K., Keller, J.M., Schubert, D., et al. (2020). m.3243A > G-Induced Mitochondrial Dysfunction Impairs Human Neuronal Development and Reduces Neuronal Network Activity and Synchronicity. Cell Rep 31, 107538. 10.1016/j.celrep.2020.107538.

72. Linda, K., Lewerissa, E.I., Verboven, A.H.A., Gabriele, M., Frega, M., Klein Gunnewiek, T.M., Devilee, L., Ulferts, E., Hommersom, M., Oudakker, A., et al. (2022). Imbalanced autophagy causes synaptic deficits in a human model for neurodevelopmental disorders. Autophagy 18, 423–442. 10.1080/15548627.2021.1936777.

73. Mossink, B., Verboven, A.H.A., van Hugte, E.J.H., Klein Gunnewiek, T.M., Parodi, G., Linda, K., Schoenmaker, C., Kleefstra, T., Kozicz, T., van Bokhoven, H., et al. (2021). Human neuronal networks on micro-electrode arrays are a highly robust tool to study disease-specific genotype-phenotype correlations in vitro. Stem Cell Reports 16, 2182–2196. 10.1016/j.stemcr.2021.07.001.

74. Zeiss, C.J. (2021). Comparative Milestones in Rodent and Human Postnatal Central Nervous System Development. Toxicol Pathol 49, 1368–1373. 10.1177/01926233211046933.

75. Marchetto, M.C., Hrvoj-Mihic, B., Kerman, B.E., Yu, D.X., Vadodaria, K.C., Linker, S.B., Narvaiza, I., Santos, R., Denli, A.M., Mendes, A.P., et al. (2019). Species-specific maturation profiles of human, chimpanzee and bonobo neural cells. Elife 8. 10.7554/eLife.37527.

76. Benito-Kwiecinski, S., Giandomenico, S.L., Sutcliffe, M., Riis, E.S., Freire-Pritchett, P., Kelava, I., Wunderlich, S., Martin, U., Wray, G.A., McDole, K., and Lancaster, M.A. (2021). An early cell shape transition drives evolutionary expansion of the human forebrain. Cell 184, 2084–2102 e2019. 10.1016/j.cell.2021.02.050.

77. Gomez-Robles, A., Nicolaou, C., Smaers, J.B., and Sherwood, C.C. (2024). The evolution of human altriciality and brain development in comparative context. Nat Ecol Evol 8, 133–146. 10.1038/s41559-023-02253-z.

78. Charvet, C.J., and Finlay, B.L. (2018). Comparing Adult Hippocampal Neurogenesis Across Species: Translating Time to Predict the Tempo in Humans. Front Neurosci 12, 706. 10.3389/fnins.2018.00706.

79. Cieslar-Pobuda, A., Knoflach, V., Ringh, M.V., Stark, J., Likus, W., Siemianowicz, K., Ghavami, S., Hudecki, A., Green, J.L., and Los, M.J. (2017). Transdifferentiation and reprogramming: Overview of the processes, their similarities and differences. Biochim Biophys Acta Mol Cell Res 1864, 1359–1369. 10.1016/j.bbamcr.2017.04.017.

80. Schlaeger, T.M., Daheron, L., Brickler, T.R., Entwisle, S., Chan, K., Cianci, A., DeVine, A., Ettenger, A., Fitzgerald, K., Godfrey, M., et al. (2015). A comparison of non-integrating reprogramming methods. Nat Biotechnol 33, 58–63. 10.1038/nbt.3070.

81. Hu, K. (2014). Vectorology and factor delivery in induced pluripotent stem cell reprogramming. Stem Cells Dev 23, 1301–1315. 10.1089/scd.2013.0621.

82. Trevisan, M., Desole, G., Costanzi, G., Lavezzo, E., Palu, G., and Barzon, L. (2017). Reprogramming Methods Do Not Affect Gene Expression Profile of Human Induced Pluripotent Stem Cells. Int J Mol Sci 18. 10.3390/ijms18010206.

83. Oberheim, N.A., Wang, X., Goldman, S., and Nedergaard, M. (2006). Astrocytic complexity distinguishes the human brain. Trends Neurosci 29, 547–553. 10.1016/j.tins.2006.08.004.

84. Drago, F., Lombardi, M., Prada, I., Gabrielli, M., Joshi, P., Cojoc, D., Franck, J., Fournier, I., Vizioli, J., and Verderio, C. (2017). ATP Modifies the Proteome of Extracellular Vesicles Released by Microglia and Influences Their Action on Astrocytes. Front Pharmacol 8, 910. 10.3389/fphar.2017.00910.

85. Matejuk, A., and Ransohoff, R.M. (2020). Crosstalk Between Astrocytes and Microglia: An Overview. Front Immunol 11, 1416. 10.3389/fimmu.2020.01416.

86. Nakanishi, M., Niidome, T., Matsuda, S., Akaike, A., Kihara, T., and Sugimoto, H. (2007). Microglia-derived interleukin-6 and leukaemia inhibitory factor promote astrocytic differentiation of neural stem/progenitor cells. Eur J Neurosci 25, 649–658. 10.1111/j.1460-9568.2007.05309.x.

87. Nemes-Baran, A.D., White, D.R., and DeSilva, T.M. (2020). Fractalkine-Dependent Microglial Pruning of Viable Oligodendrocyte Progenitor Cells Regulates Myelination. Cell Rep 32, 108047. 10.1016/j.celrep.2020.108047.

88. McNamara, N.B., Munro, D.A.D., Bestard-Cuche, N., Uyeda, A., Bogie, J.F.J., Hoffmann, A., Holloway, R.K., Molina-Gonzalez, I., Askew, K.E., Mitchell, S., et al. (2023). Microglia regulate central nervous system myelin growth and integrity. Nature 613, 120–129. 10.1038/s41586-022-05534-y.

89. Joseph, G., Orme, R.P., Kyriacou, T., Fricker, R.A., and Roach, P. (2021). Effects of Surface Chemistry Interaction on Primary Neural Stem Cell Neurosphere Responses. ACS Omega 6, 19901–19910. 10.1021/acsomega.1c02796.

90. Noora Räsänen, J.T., Marja Koskuvi, Šárka Lehtonen, Nelli Jalkanen, Nelli Karmila, Isabelle Weert, Olli Vaurio, Ilkka Ojansuu, Markku Lähteenvuo, Olli Pietiläinen, Jari Koistinaho (2024). Astrocytes regulate neuronal network burst frequency through NMDA receptors species- and donor-specifically. BioRxiv. 10.1101/2023.12.09.570906.

