## Supplementary Figures for "Navigating Human Astrocyte Differentiation: Direct and Rapid one-step Differentiation of Induced Pluripotent Stem Cells to Functional Astrocytes Supporting Neuronal Network development"

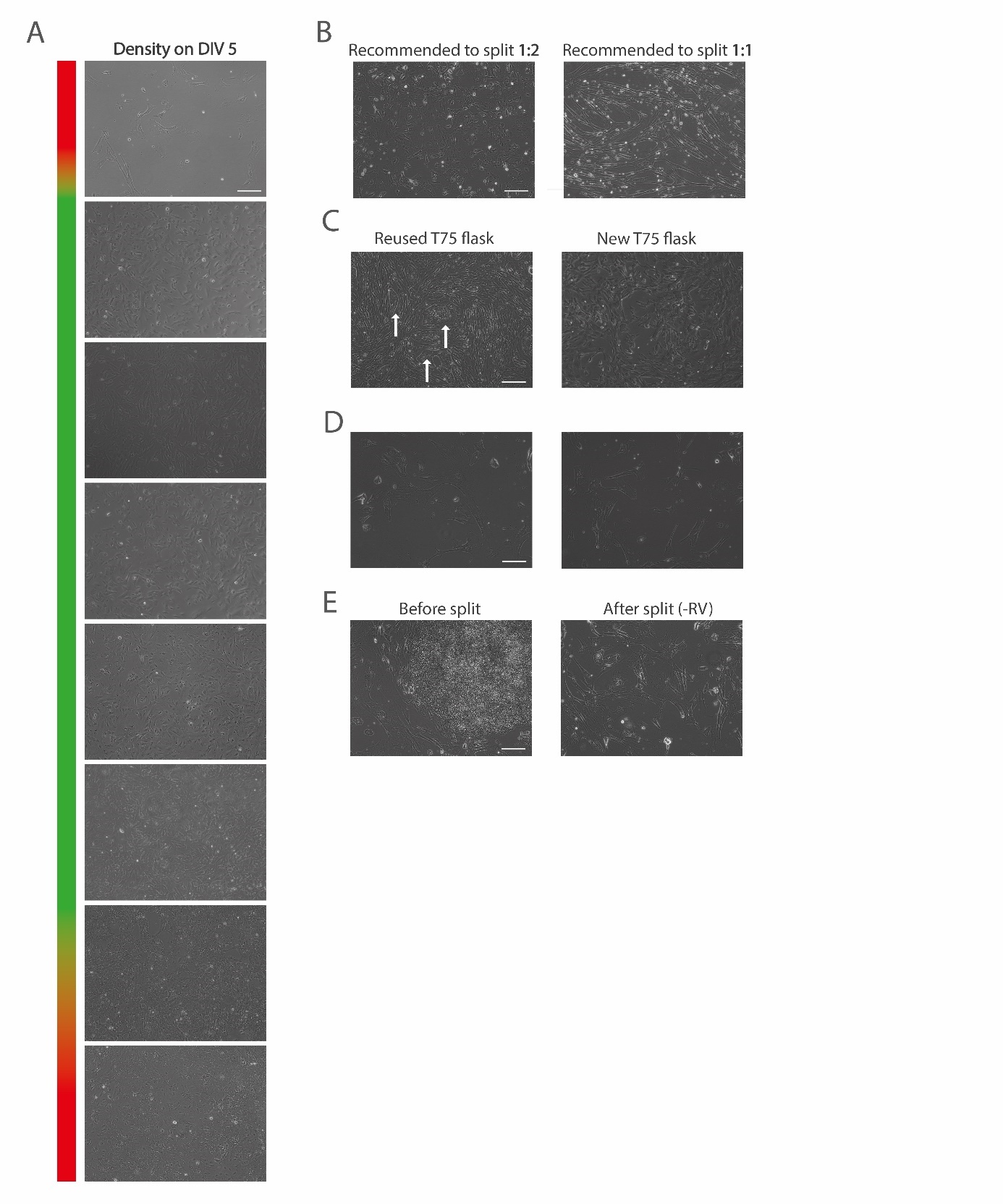

**Supplementary Figure 1. (A)** Brightfield images of different hiPSC-astrocyte cultures at DIV 5, ranging from (too) low to (too) high astrocyte densities at DIV 5. Color bar is indicative of acceptable and unacceptable astrocyte densities. All pictures were taken at the same magnification (scale bar = 100 µm). **(B)** Brightfield images showing astrocytes cultures that are recommended to be split either 1:1 or 1:2 ratio in order to rescue the differentiation and/or proliferation of these cultures. All pictures were taken at the same magnification (scale bar = 100 µm). **(C)** Brightfield images highlighting the importance of not reusing T75 flasks for splitting, but instead using new flaks to prevent insufficient astrocyte growth and the culture of hiPSC-like colonies (indicated by white arrows). All pictures were taken at the same magnification (scale bar = 100 µm). **(D)** Brightfield images of two example astrocyte cultures that showed a rapidly decreased proliferation rate within the first week(s). All pictures were taken at the same magnification (scale bar = 100 µm). **(E)** The left brightfield image shows an astrocyte culture with so-called delayed astrocyte differentiation, indicated be a remaining hiPSC-colony, before it has been split. The right brightfield image shows to same astrocyte culture after splitting without Revitacell^TM^, which improves astrocyte differentiation and removes the remaining hiPSC-colonies. All pictures were taken at the same magnification (scale bar = 100 µm).

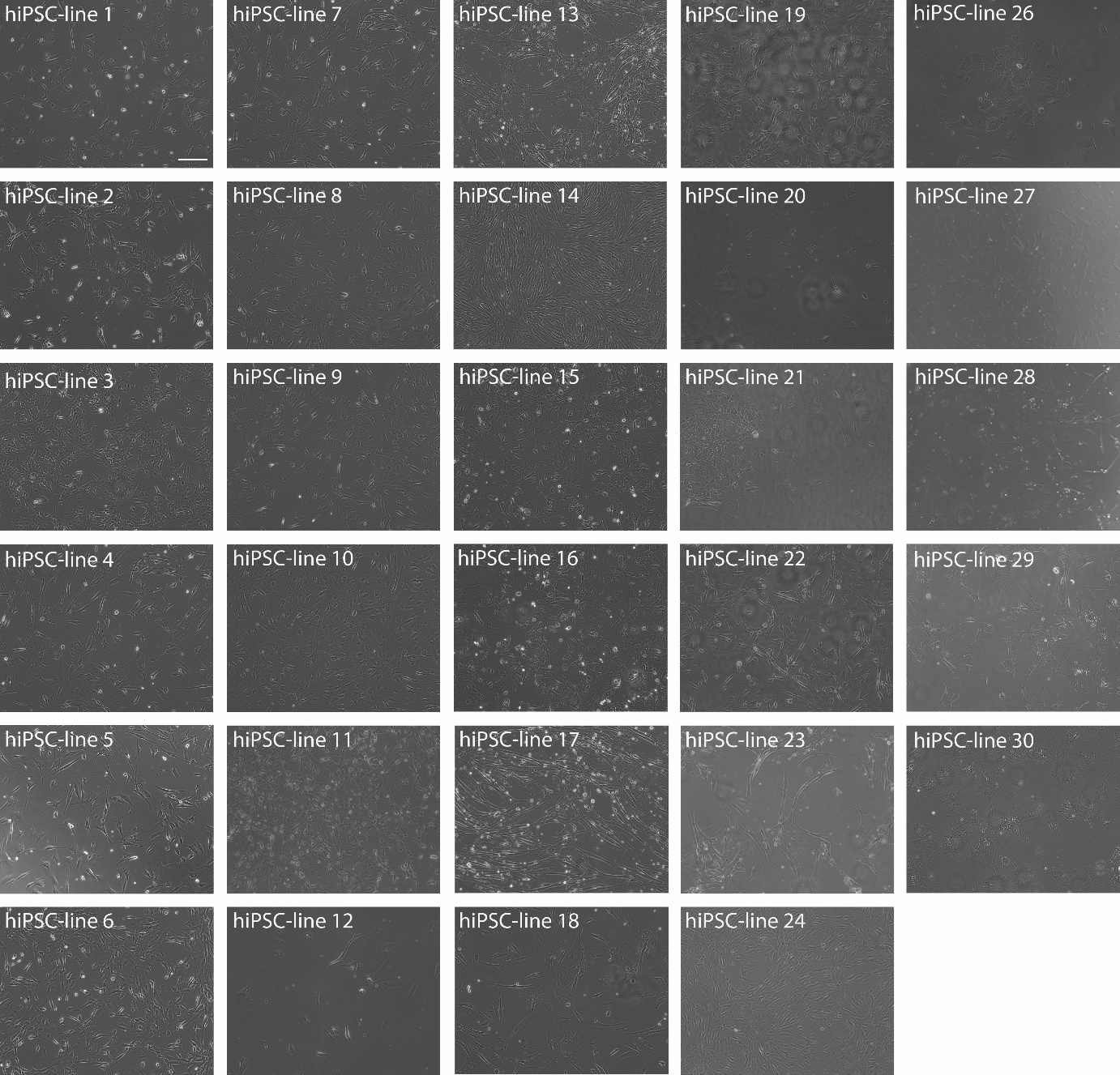

**Supplementary Figure 2.** Brightfield images of twenty-nine (from all thirty) different hiPSC-astrocyte cultures, which were cultured across different batches and by several different researchers, highlighting morphological differences between the different astrocyte cultures. The astrocytes are around 2.5-4 weeks old. All pictures were taken at the same magnification (scale bar = 100 µm).

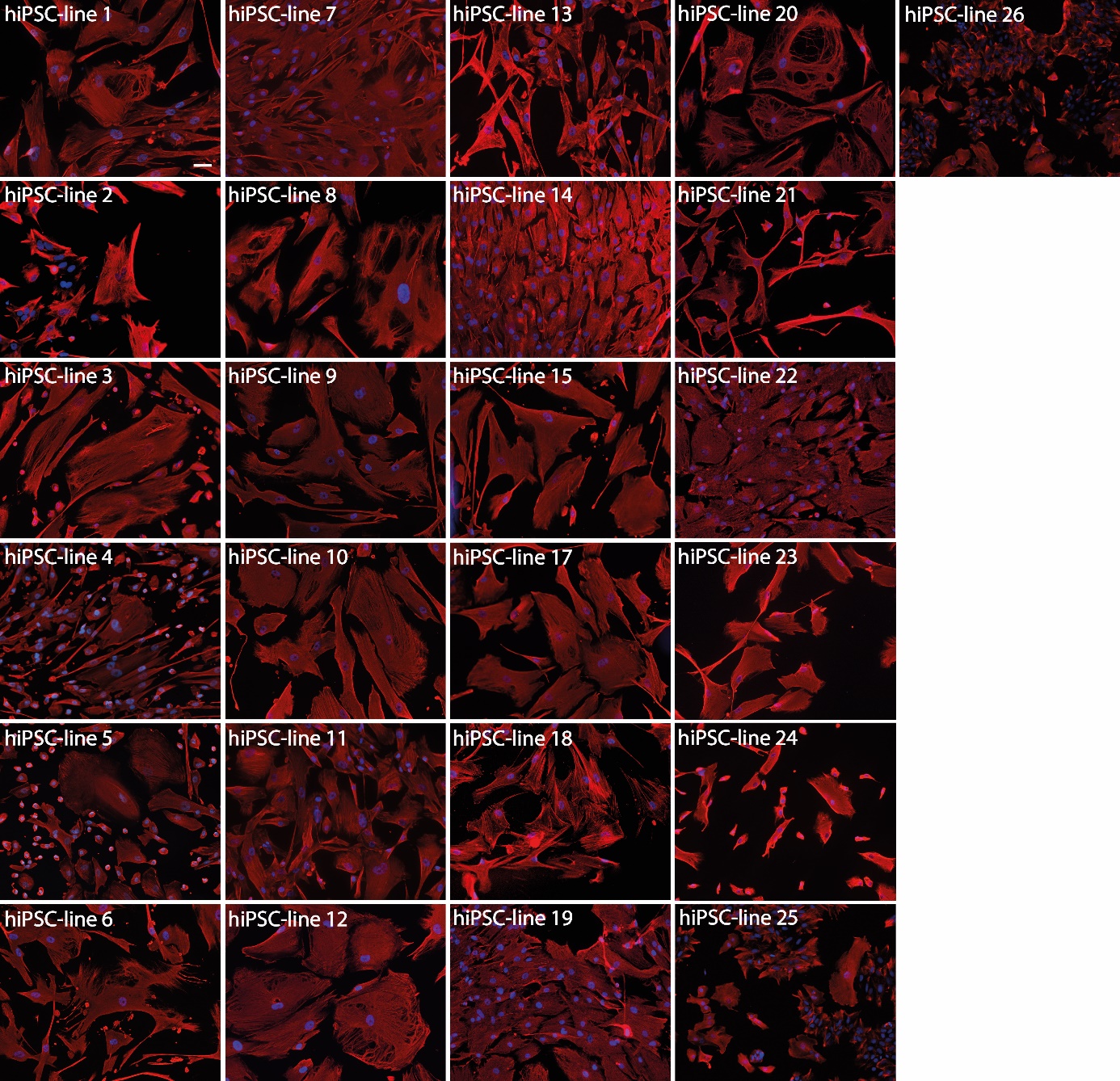

**Supplementary Figure 3.** Immunostaining of Vimentin (red) by twenty-five from the twenty-eight different hiPSC-astrocyte cultures, which were cultured across different batches and by several different researchers, highlighting morphological differences between the different astrocyte cultures. The astrocytes are around 5-6 weeks old. All pictures were taken at the same magnification (scale bar = 50 µm).

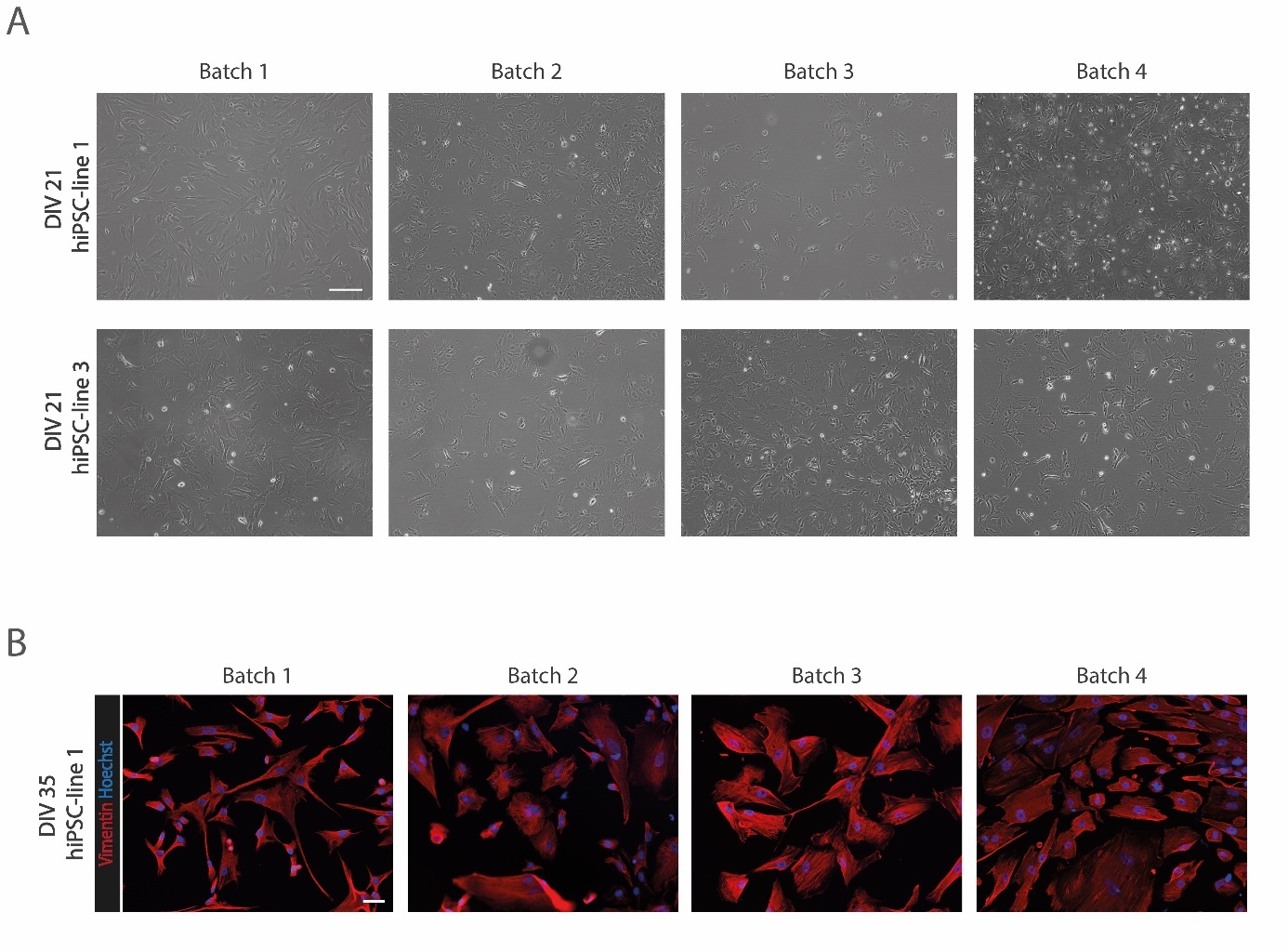

**Supplementary Figure 4. (A)** Brightfield images of DIV 21 astrocytes derived from hiPSC-line 1 and hiPSC-line 3 across four different batches, highlighting minimal batch-to-batch variability based on morphology. All pictures were taken at the same magnification (scale bar = 100 µm). **(B)** Immunostaining of Vimentin (red) by DIV 35 astrocytes derived from hiPSC-line 1 across four different batches, highlighting minimal batch-to-batch variability based on morphology and vimentin expression. All pictures were taken at the same magnification (scale bar = 50 µm).

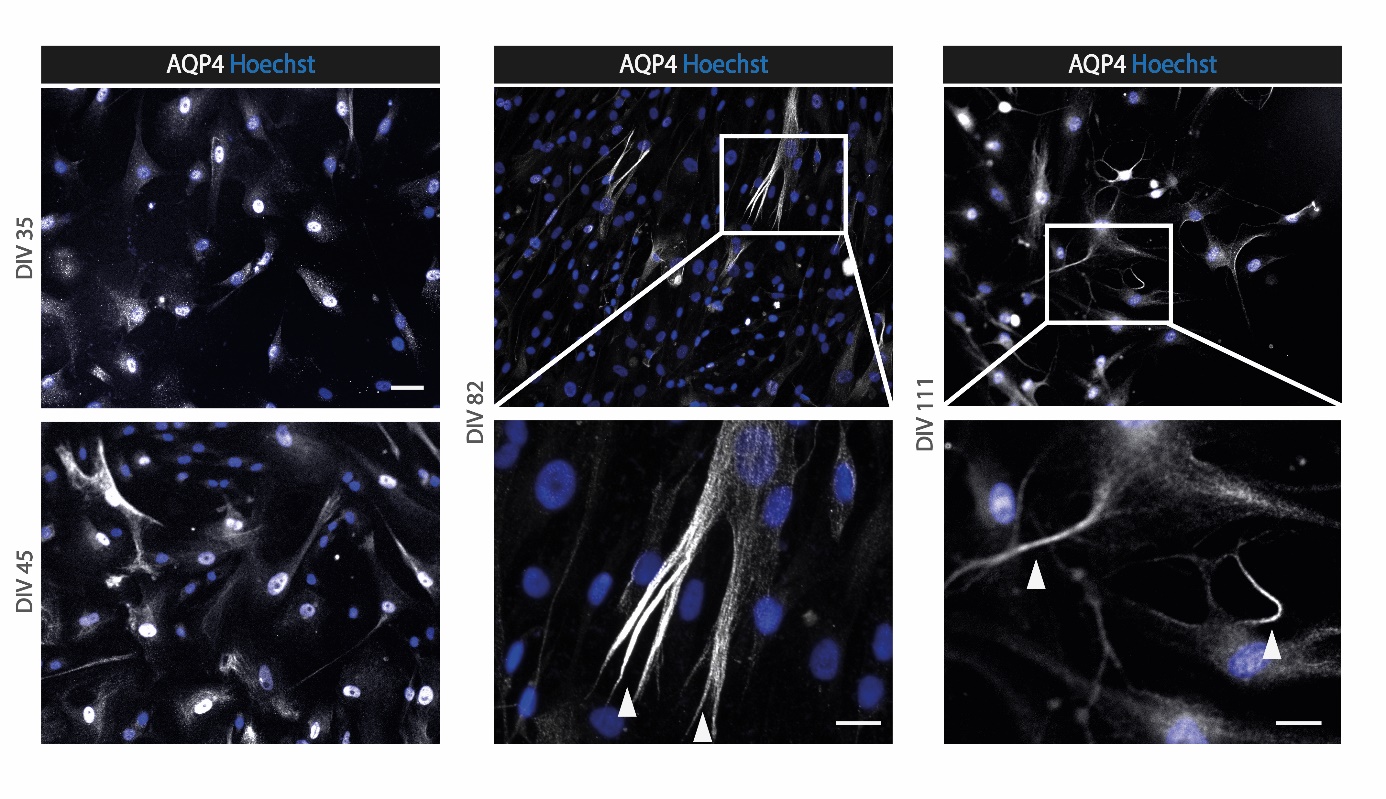

**Supplementary Figure 5.** Expression of AQP4 (white) by immunolabeling in astrocytes from hiPSC-line 1 at DIV 35, 45, 82 and DIV 111. The images show that AQP4 is expressed throughout the whole astrocyte cell body at early stages, while later during differentiation the expression of AQP4 decreases as it localises towards the astrocytic end feet (indicated by the white arrows). All pictures were taken at the same magnification (scale bar = 50 µm or 10 µm in the zoomed image).

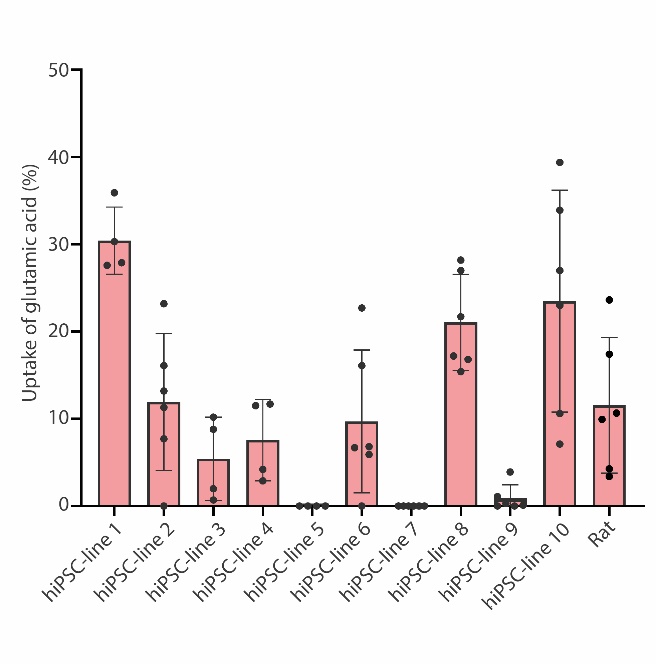

**Supplementary Figure 6.** The percentage of glutamic acid uptake by monoculture hiPSC-astrocytes was measured for ten different hiPSC-astrocyte cultures and one rat astrocyte culture.

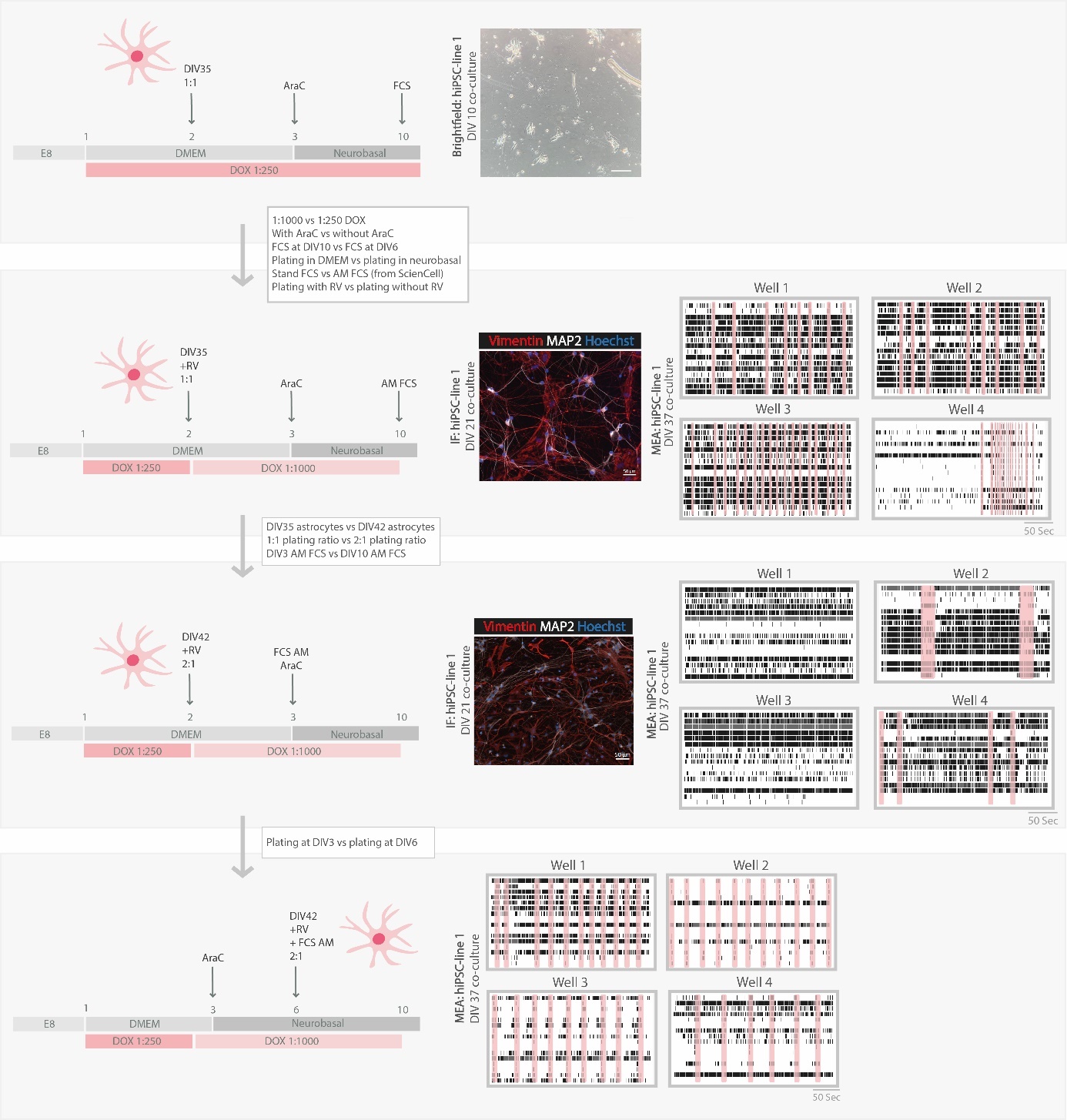

**Supplementary Figure 7.** Schematic representation of the optimization of the co-culture is depicted. As starting point the “standard” co-culture protocol was tested. Brightfield imaging, immunofluorescent imaging and/or MEA were used as the main readout to assess the quality of the co-cultures. Upon testing many adjusted versions of the protocol, the bottom protocol is considered most optimal and reproducible. All pictures were taken at the same magnification (scale bar = 50 µm).
